# Quantifying the Benefit Offered by Transcript Assembly on Single-Molecule Long Reads

**DOI:** 10.1101/632703

**Authors:** Laura H. Tung, Mingfu Shao, Carl Kingsford

## Abstract

Third-generation sequencing technologies benefit transcriptome analysis by generating longer sequencing reads. However, not all single-molecule long reads represent full transcripts due to incomplete cDNA synthesis and the sequencing length limit of the platform. This drives a need for long read transcript assembly. We quantify the benefit that can be achieved by using a transcript assembler on long reads. Adding long-read-specific algorithms, we evolved Scallop to make Scallop-LR, a long-read transcript assembler, to handle the computational challenges arising from long read lengths and high error rates. Analyzing 26 SRA PacBio datasets using Scallop-LR, Iso-Seq Analysis, and StringTie, we quantified the amount by which assembly improved Iso-Seq results. Through combined evaluation methods, we found that Scallop-LR identifies 2100–4000 more (for 18 human datasets) or 1100–2200 more (for eight mouse datasets) known transcripts than Iso-Seq Analysis, which does not do assembly. Further, Scallop-LR finds 2.4–4.4 times more potentially novel isoforms than Iso-Seq Analysis for the human and mouse datasets. StringTie also identifies more transcripts than Iso-Seq Analysis. Adding long-read-specific optimizations in Scallop-LR increases the numbers of predicted known transcripts and potentially novel isoforms for the human transcriptome compared to several recent short-read assemblers (e.g. StringTie). Our findings indicate that transcript assembly by Scallop-LR can reveal a more complete human transcriptome.

## 1 Introduction

More than 95% of human genes are alternatively spliced to generate multiple isoforms (Pan *et al.*, 2008). Gene regulation through alternative splicing can create different functions for a single gene and increase protein-coding capacity and proteomic diversity. Thus, studying the full transcriptome is crucial to understanding the functionality of the genome. In the past decade, high-throughput, short-read sequencing technologies have become powerful tools for the characterization and quantification of the transcriptome. However, due to limited read lengths, identifying full-length transcripts from short reads and assembling all spliced RNAs within a transcriptome remain challenging problems. In recent years, third-generation sequencing technologies offered by Pacific Biosciences (PacBio) and Oxford Nanopore Technologies (ONT) produce sequences of full cDNA or RNA molecules, promising to improve isoform identification and reducing ambiguity in mapping reads (Cho *et al.*, 2014). Long reads offer various benefits such as covering the entire molecule in the majority of cases and determining the allele from which the RNA molecule originated by identifying single nucleotide variations (SNVs) affecting each single RNA molecule (Tilgner *et al.*, 2014). Long reads are also able to capture gene structures accurately without annotation and identify novel splice patterns that are not found by short reads (Cho *et al.*, 2014). Long reads have been used for genome assembly and can be used to identify functional elements in genomes that are missed by short-read sequencing (Shi *et al.*, 2016; Koren *et al.*, 2017; Zimin *et al.*, 2013). Hybrid sequencing combining long reads and short reads can improve isoform identification and transcriptome characterization (Au *et al.*, 2013; Weirather *et al.*, 2015). Long reads are also useful in identifying novel long non-coding RNAs and fusion transcripts (Wang *et al.*, 2016) and in studying specific disease determinant genes (Tseng *et al.*, 2017).

A main challenge associated with long-read technologies is high error rates. PacBio produces reads with average lengths up to 30 kb, and its error rate for “subreads” (raw reads, which are original lower quality reads as opposed to consensus reads) is ~10–20%. Continuous Long Read (CLR) is the original polymerase read (by reading a template with the DNA polymerase), and subreads are sequences generated by splitting the CLR by the adapters (a full-pass subread is flanked on both ends by adapters). However, PacBio’s “ROI” (“Read of Insert”, consensus reads) displays a higher quality than subreads. Circular Consensus Sequence (CCS) reads are a type of ROI and are generated by collapsing multiple subreads when ≥ 2 full-pass subreads are present. ONT produces longer reads with even higher error rates (error rates for “1D” raw reads: > 25%; error rates for “2D” consensus reads: 12–20%) (Križanović *et al.*, 2018). Error-correction methods using short reads (such as the error correction tool LSC (Au *et al.*, 2012)) have been created to correct the high rate of errors in long reads; however, error correction may create artifacts so that the corrected long reads may no longer be true single-molecule reads (Sharon *et al.*, 2013).

We focus on transcript assembly of long reads, aiming to discover more novel isoforms. Although it is often thought that long reads are full-length transcripts and isoforms with no assembly required^1^, in fact the success rate of the sequenced cDNA molecules containing all splice sites of the original transcripts depends on the completeness of cDNA synthesis (Sharon *et al.*, 2013). Sharon *et al.* (2013) found that a CCS read could correspond to an incomplete transcript as a consequence of incomplete cDNA synthesis, although a CCS read represents the full cDNA molecule. They found that, in their experiment, for transcripts > 2.5 kb, full-length reads that represent the original transcripts are less likely to be observed than those for transcripts < 2.5 kb. Tilgner *et al.* (2014) also found that, in their experiment, reads representing all splice sites of the original transcripts are more likely to be observed for transcripts ≤ 3 kb. The cDNA synthesis methods impose limitations on long reads (Kuosmanen *et al.*, 2016) even though with increasing performance the sequencing technologies can be capable of sequencing long full-length transcripts. In addition, long reads may still be limited by the sequencing length limit of the platform (Rhoads and Au, 2015). Thus, incomplete cDNA synthesis plus the sequencing length limit could cause PacBio’s consensus long reads to miss a substantial number of true transcripts (Rhoads and Au, 2015), especially longer transcripts. This suggests that the transcript assembly of long reads is still needed, since it is possible that those CCS reads corresponding to incomplete transcripts could be assembled together to recover the original full transcripts.

Long read lengths and high error rates pose computational challenges to transcript assembly. No published transcript assembler has been adapted and systematically tested on the challenges of long-read transcript assembly yet. Aiming to handle these challenges, we developed a longread transcript assembler called Scallop-LR, evolved from Scallop, an accurate short-read transcript assembler (Shao and Kingsford, 2017). Scallop-LR is designed for PacBio long reads. Scallop-LR’s algorithms are tailored to long-read technologies, dealing with the long read lengths and high error rates as well as taking advantage of long-read-specific features such as the read boundary information to construct more accurate splice graphs. A post-assembly clustering algorithm is also added in Scallop-LR to reduce false negatives.

We analyzed 26 long-read datasets from NIH’s Sequence Read Archive (SRA) (Leinonen *et al.*, 2011) with Scallop-LR, Iso-Seq Analysis^2^, and StringTie (Pertea *et al.*, 2015; 2016). Iso-Seq Analysis, also known as Iso-Seq informatics pipeline, is a software system developed by PacBio that takes subreads as input and outputs polished isoforms (transcripts) through collapsing, clustering, consensus calling, etc. Iso-Seq Analysis does not perform assembly per se. The clustering algorithm in Iso-Seq Analysis clusters reads based on their isoform of origin. An algorithm that clusters long reads based on their gene family of origin was recently proposed (Sahlin and Medvedev, 2019). StringTie was originally designed as a short-read transcript assembler but can also assemble long reads. StringTie outperforms many leading short-read transcript assemblers (Pertea *et al.*, 2015).

Through combined evaluation methods, we demonstrate that Scallop-LR is able to find more known transcripts and novel isoforms that are missed by Iso-Seq Analysis. We show that Scallop-LR can identify 2100–4000 more known transcripts (in each of 18 human datasets) or 1100–2200 more known transcripts (in each of eight mouse datasets) than Iso-Seq Analysis. The sensitivity of Scallop-LR is 1.33–1.71 times higher (for the human datasets) or 1.43–1.72 times higher (for the mouse datasets) than that of Iso-Seq Analysis. Scallop-LR also finds 2.53–4.23 times more (for the human datasets) or 2.38–4.36 times more (for the mouse datasets) potential novel isoforms than Iso-Seq Analysis. Further, Scallop-LR assembles 950–3770 more known transcripts and 1.37–2.47 times more potential novel isoforms than StringTie, and has 1.14–1.42 times higher sensitivity than StringTie for the human datasets.

## 2 Methods

### 2.1 Scallop-LR Algorithms for Long-Read Transcript Assembly

Scallop-LR is a reference-based transcript assembler that follows the standard paradigm of alignment and splice graphs but has a computational formulation dealing with “phasing paths.” “Phasing paths” are a set of paths that carry the phasing information derived from the reads spanning more than two exons. The reads are first aligned to a reference genome and the alignments are transformed into splice graphs, in which vertices are exons, edges are splice junctions, the coverage of exon is taken as the vertex weight, and the abundance of splice junction is used as the edge weight. We decompose the splice graph to infer a small number of paths (i.e. predicted transcripts) that cover the topology and fit the weights of the splice graph.

#### 2.1.1 Scallop-LR represents long reads as long phasing paths, preserved in assembly

Unlike short reads, most long reads span more than two exons. Thus, if the multi-exon paths of long reads are broken when decomposing splice graphs (which is more likely to occur since the majority of long reads span large numbers of exons), many long reads would not be correctly covered by assembled transcripts. Thus, Scallop-LR represents long reads as long phasing paths and preserves phasing paths in assembly. This is particularly important since we want every phasing path (and thus every long read) to be covered by some transcript so that the assembly can represent the original mRNAs. Scallop-LR adapted the phasing-path preservation algorithm from Scallop when decomposing splice graphs into transcripts. The Scallop algorithm uses an iterative strategy to gradually decompose the splice graph while achieving three objectives simultaneously:

a. Preserving all phasing paths in assembled transcripts when decomposing the splice graph;
b. Minimizing the read coverage deviation using linear programming;
c. Minimizing the number of predicted transcripts by reducing an upper bound on the number of required paths.

By representing long reads as long phasing paths, Scallop-LR makes full use of the information in long reads through phasing-path preservation, so that assembled transcripts can best represent the input long reads.

#### 2.1.2 Added Scallop-LR algorithms

To improve the long-read assembly accuracy, Scallop-LR extracts the boundary information from long reads and identifies transcript boundaries to build a more accurate splice graph. In single-molecule sequencing, there are two types of long reads produced: full-length reads and non-full-length reads. Full-length reads are the reads that have a 5’ primer, 3’ primer, and polyA tail, which are the reads that represent full-length transcripts they originated from. Non-full-length reads do not represent full-length transcripts. We further classify non-full-length reads into two types: non-full-length boundary reads and non-full-length internal reads. Non-full-length boundary reads are the reads that either have a 5’ primer but not the 3’ primer, or have a 3’ primer but not the 5’ primer (i.e. reads that come from either the 5’ or 3’ end but do not reach the other end). Non-full-length internal reads are the reads that have neither of the 5’ primer and 3’ primer (i.e. reads that do not come from either end). Scallop-LR treats non-full-length internal reads like short reads when constructing the splice graph.

We refer to non-full-length boundary reads (with one side boundary) and full-length reads (with two side boundaries) as “boundary reads” for the side they have a boundary. We use the *Classify* tool in Iso-Seq Analysis to obtain full-length and non-full-length CCS reads. The Scallop-LR algorithm extracts the boundary information of each read from the Classify results and uses it to deduce starting/ending boundaries in the splice graph. Specifically, when there are a certain number of boundary reads within a region (the default minimum number is 3), the algorithm defines it as a starting or ending boundary:

> Suppose there are some 5’ end boundary reads aligned to the genome at positions [*a* + *δ*_1_, *x*_1_], [*a* + *δ*_2_, *x*_2_], [*a* + *δ*_3_, *x*_3_], etc., where *|δ*_1_*|*, *|δ*_2_*|*, *|δ*_3_*|*,… are within a predefined allowance of difference for matching positions, then this is a signal that position “***a***” corresponds to a starting position of a transcript. Thus, in the splice graph, we add an edge connecting the source ***s*** to the vertex corresponding to the region [*a, c*].

Similarly,

> Suppose there are some 3’ end boundary reads aligned to the genome at positions [*x*_1_, *b* + *δ*_1_], [*x*_2_, *b* + *δ*_2_], [*x*_3_, *b* + *δ*_3_], etc., where *|δ*_1_*|*, *|δ*_2_*|*, *|δ*_3_*|*,… are within a predefined allowance of difference for matching positions, then this is a signal that position “***b***” corresponds to an ending position of a transcript. Thus, in the splice graph, we add an edge connecting the vertex corresponding to the region [*d, b*] to the target ***t***.

This is for the forward strand. For the reverse strand, the situation is opposite.

To increase the precision of long-read assembly, Scallop-LR uses a post-assembly clustering algorithm to reduce the false negatives in the final predicted transcripts. For transcripts with very similar splice positions, the algorithm clusters them into a single transcript. The “very similar splice positions” means (a) these transcripts have the same number of splice positions; (b) for each splice position, their position differences are within a predefined allowance (the default allowance is 10 bp; the allowance can be set in a parameter). This allowance is for the sum of the difference (absolute value) of starting position and the difference of ending position for a splice position. This clustering collapses “nearly redundant” transcripts and thus increases the precision of assembly.

The Scallop-LR algorithm deals with the high error rates in long reads when building the splice graph. When identifying splice positions from long-read alignments during the construction of the splice graph, the algorithm takes into account that a single insertion or deletion in the middle of the alignment between introns may be caused by errors in long reads and treats the single insertion or deletion between introns as alignment match when determining the splice positions.

### 2.2 Combined Evaluation Methods

We use multiple transcript evaluation methods to examine the quality of predicted transcripts from transcript assemblers (i.e. Scallop-LR and StringTie) and Iso-Seq Analysis. The combined evaluation methods allow us to assess predicted transcripts using various metrics as well as cross-verify the findings obtained from different methods.

Gffcompare^3^ is used to identify correctly predicted transcripts and the resulting sensitivity and precision by comparing the intron chains of predicted transcripts to the reference annotation for matching intron-exon structures. A correctly predicted known transcript has an exact intron-chain matching with a reference transcript. Sensitivity is the ratio of the number of correctly predicted known transcripts over the total number of known transcripts, and precision is the ratio of the number of correctly predicted known transcripts over the total number of predicted transcripts. We generate the precision-recall curve (PR-curve) based on the results of Gffcompare by varying the set of predicted transcripts sorted with coverage, and compute the metric PR-AUC (area under the PR-curve) which measures the overall performance. Gffcompare also reports “potential novel isoforms” that are predicted transcripts sharing at least one splice junction with reference transcripts, though this criterion for potential novel isoforms is weak when transcripts contain many splice junctions.

To further examine novel isoforms, we use the evaluation method SQANTI (Tardaguila *et al.*, 2018) that classifies novel isoforms into Novel in Catalog (NIC) and Novel Not in Catalog (NNC). A transcript classified as NIC either contains new combinations of known splice junctions or contains novel splice junctions formed from known donors and acceptors. NNC contains novel splice junctions formed from novel donors and/or novel acceptors. The criterion for NIC is stronger compared with that of potential novel isoforms in Gffcompare, and we conjecture that NICs may be more likely to be true novel isoforms than wrongly assembled transcripts. SQANTI also reports Full Splice Match (FSM) that is a predicted transcript matching a reference transcript at all splice junctions, and Incomplete Splice Match (ISM) that is a predicted transcript matching consecutive, but not all, splice junctions of a reference transcript.

Gffcompare and SQANTI report transcripts that fully match, partially match, or do not match reference transcripts, but do not report how many transcripts, for example, have 75–95% or 50–75% of bases matching a reference transcript. These ranges of matched fractions would give us a more detailed view of the overall quality of assembly. Thus, we use rnaQUAST (Bushmanova *et al.*, 2016) that measures the fraction of a predicted transcript matching a reference transcript. rnaQUAST maps predicted transcripts sequences to the reference genome using GMAP (Wu and Watanabe, 2005) and matches the alignments to the reference transcripts’ coordinates from the gene annotation database. rnaQUAST measures the fraction of a reference transcript that is covered by a single predicted transcript, and the fraction of a predicted transcript that matches a reference transcript. Based on the results of rnaQUAST, we compute the distribution of predicted transcripts in different ranges of fractions matching reference transcripts, and the distribution of reference transcripts in different ranges of fractions covered by predicted transcripts. rnaQUAST also reports unaligned transcripts (transcripts without any significant alignments), misassembled transcripts (transcripts that have discordant best-scored alignments, i.e. partial alignments that are mapped to different strands, different chromosomes, in reverse order, or too far away), and unannotated transcripts (predicted transcripts that do not cover any reference transcript).

We use Transrate (Smith-Unna *et al.*, 2016) for sequence-based evaluation to obtain statistics of predicted transcripts such as the minimum, maximum, and mean lengths, the number of bases in the assembly, numbers of transcripts in different size ranges, etc.

The reference annotations we use in Gffcompare, rnaQUAST, and SQANTI are Ensembl *Homo sapiens* GRCh38.90 and *Mus musculus* GRCm38.92. The reference genomes we use are Ensembl GRCh38 for human and GRCm38 for mouse when running rnaQUAST and SQANTI or aligning long reads to the genome (Section 2.4).

### 2.3 Data Acquisition and Preprocessing

We obtained PacBio datasets for *Homo sapiens* and *Mus musculus* from SRA (Leinonen *et al.*, 2011; Shi *et al.*, 2016; Komor *et al.*, 2017; O’Grady *et al.*, 2016; Seo *et al.*, 2016; Hughes *et al.*, 2010). In most of the PacBio datasets in SRA, one BioSample has multiple SRA Runs because the experimenters used multiple “movies” to increase the coverage so that low-abundance, long isoforms can be captured in analysis. The experimenters also used a size selection sequencing strategy, and thus different SRA Runs are designated for different size ranges. Therefore, we use one BioSample instead of one SRA Run to represent one dataset in our analysis, and we merge multiple SRA Runs that belong to the same BioSample into that dataset. (See Supplementary Materials section 1 about “movies” and size selection strategy).

We collected the SRA PacBio datasets that meet the following conditions: (a) The datasets should be transcriptomic and use the cDNA library preparation. (b) The datasets should have the *hdf5* raw data uploaded. This is because if using *fastq-dump* in SRA Toolkit to extract the sequences from SRA, the output sequences lose the original PacBio sequence names even using the sequence-name preserving option. The original PacBio sequence name is critical since it contains information such as the movie, the identification of subreads or CCS reads, etc. (c) The datasets should not be “targeted sequencing” focusing on a specific gene or a small genomic region. (d) The datasets should use the Iso-Seq2 supported sequencing-chemistry combinations. (e) For a BioSample, the number of SRA Runs should be ≤ 50. This is because a huge dataset is very computationally expensive for Iso-Seq Analysis. With the above conditions, we identified and extracted 18 human datasets and eight mouse datasets—a total of 26 PacBio datasets from SRA. These 26 datasets are sequenced using RS II or RS platform, and their SRA information is in Table S9.

We convert the PacBio raw data to subreads and merge the subreads from multiple movies belonging to the same BioSample into a large dataset for analysis.

### 2.4 Analysis Workflow for Analyzing the SRA PacBio Datasets

Combining our long-read transcript assembly pipeline with the Iso-Seq Analysis pipeline (Iso-Seq2), we build an analysis workflow to analyze the SRA datasets, as shown in Figure 1.

**Figure 1:**
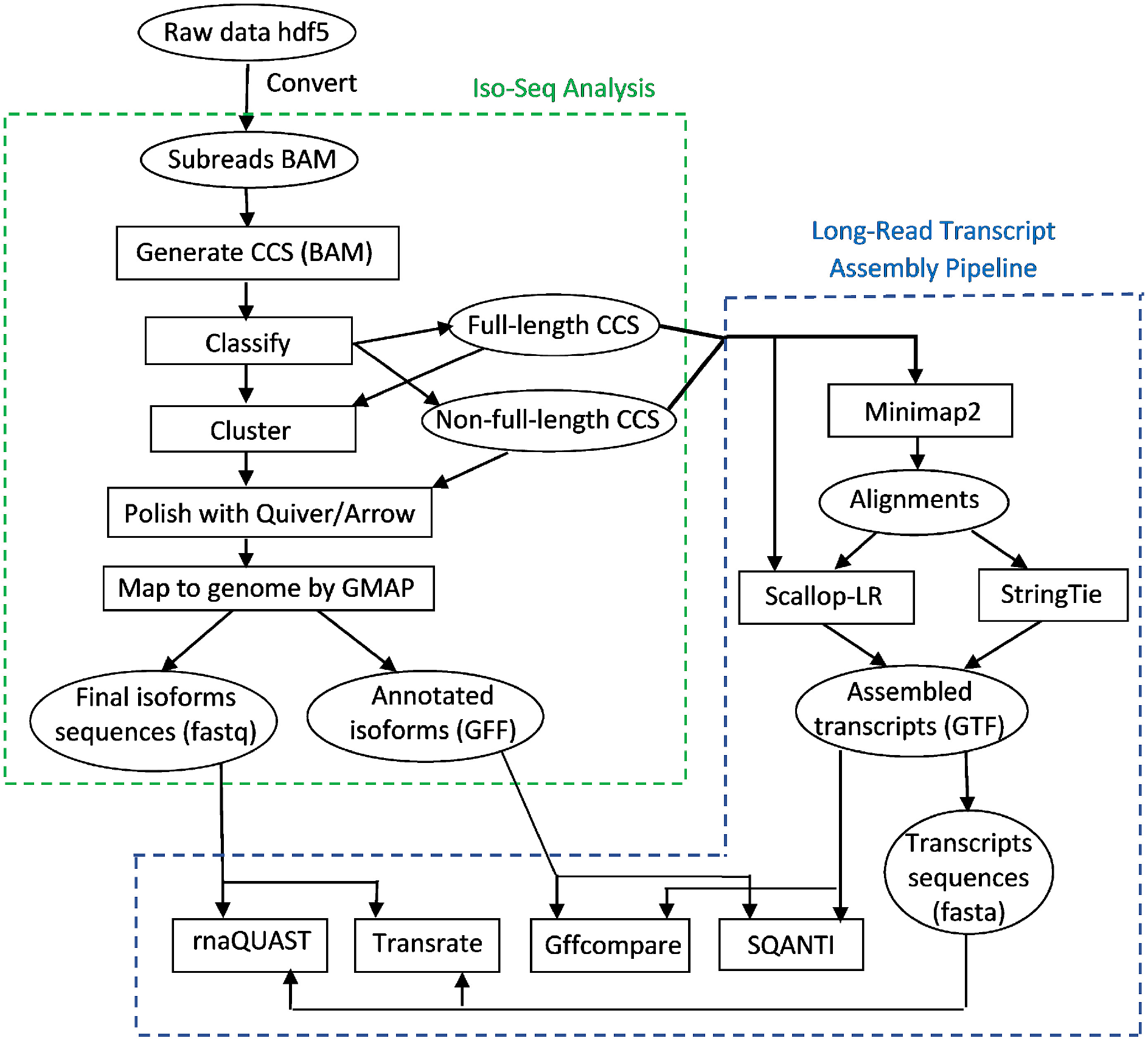
Workflow for analyzing the SRA PacBio datasets, combining the long-read transcript assembly pipeline (right) with the Iso-Seq Analysis pipeline (left).

After obtaining subreads and creating the merged dataset, we generate CCS reads from subreads. After classifying the CCS reads into full-length and non-full-length reads, the full-length CCS reads are clustered—they are run through the ICE (Iterative Clustering and Error correction) algorithm to generate clusters of isoforms. Afterwards, the non-full-length CCS reads are attributed to the clusters, and the clusters are polished using Quiver or Arrow. Quiver is an algorithm for calling accurate consensus from multiple reads, using a pair-HMM exploiting the basecalls and QV (quality values) metrics to infer the true underlying sequence^4^. Quiver is used for RS and RS II data (for data from the Sequel platform, an improved consensus model Arrow is used). Finally, the polished consensus isoforms are mapped to the genome using GMAP to remove the redundancy, and the final polished isoforms sequences and annotated isoforms are generated.

The right side of the analysis workflow in Figure 1 is our long-read transcript assembly pipeline. We chose Minimap2 (Li, 2017) and GMAP as the long-read aligners. GMAP has been shown to outperform RNA-seq aligners STAR (Dobin *et al.*, 2013), TopHat2 (Kim *et al.*, 2013), HISAT2 (Kim *et al.*, 2015), and BBMap (Bushnell, 2014) in aligning long reads (Križanović *et al.*, 2018). The recently published RNA-seq aligner Minimap2 is specifically designed for long reads. Min-imap2 outperforms GMAP, STAR, and SpAln in junction accuracy, and is 40X faster than GMAP (Li, 2017). We did a pre-assessment on the accuracy of Minimap2 vs. GMAP on a set of datasets which are either error-corrected or not error-corrected (results are not shown). Comparing the assembly results, we found that Minimap2 is more accurate than GMAP for long reads without error corrections, and Minimap2 and GMAP have nearly the same accuracy for long reads with error corrections. Thus, we use Minimap2 to align CCS reads (which are not error-corrected), while in the Iso-Seq Analysis pipeline, GMAP is used to align polished isoforms (which are error-corrected). For assembly performance comparison, we choose StringTie as a counterpart, as StringTie outperforms leading transcript assemblers Cufflinks, IsoLasso, Scripture and Traph in short-read assembly (Pertea *et al.*, 2015; 2016).

We use the full-length CCS and non-full-length CCS reads as the input of our long-read transcript assembly pipeline for Scallop-LR (v0.9.1) and StringTie (v1.3.2d) to assemble those CCS reads. We first align those CCS reads to the reference genome using Minimap2, and then the alignments are assembled by the transcript assemblers. In addition to taking the alignments as input, Scallop-LR also extracts the boundary information (see Section 2.1.2) from CCS reads.

The software versions and options used in this analysis workflow are summarized in the Supplementary Materials (section 2). The code to reproduce the analysis is available at Scallop-LR: https://github.com/Kingsford-Group/scallop/releases/tag/isoseq-v0.9.1; long-read transcript assembly analysis: https://github.com/Kingsford-Group/lrassemblyanalysis.

## 3 Results

### 3.1 Scallop-LR and StringTie predict more known transcripts than Iso-Seq Analysis

From the Gffcompare results for the human data, Scallop-LR and StringTie consistently predict more known transcripts than Iso-Seq Analysis and thus consistently have higher sensitivity than Iso-Seq Analysis. Scallop-LR finds 2100–4000 more known transcripts than Iso-Seq Analysis, and the sensitivity of Scallop-LR is 1.33–1.71 times higher than that of Iso-Seq Analysis (Figures 2 and 3, Tables S1 and S2). StringTie finds 350–1960 more known transcripts than Iso-Seq Analysis, and the sensitivity of StringTie is 1.05–1.4 times higher than that of Iso-Seq Analysis. Scallop-LR and StringTie have higher sensitivity than Iso-Seq Analysis because Scallop-LR and StringTie do assembly but Iso-Seq Analysis does not. This supports the idea that the transcript assembly of long reads is needed. Assembly is likely useful because the success level of transcriptomic long-read sequencing depends on the completeness of cDNA synthesis, and also long reads may not cover those transcripts longer than a certain length limit (Rhoads and Au, 2015).

**Figure 2:**
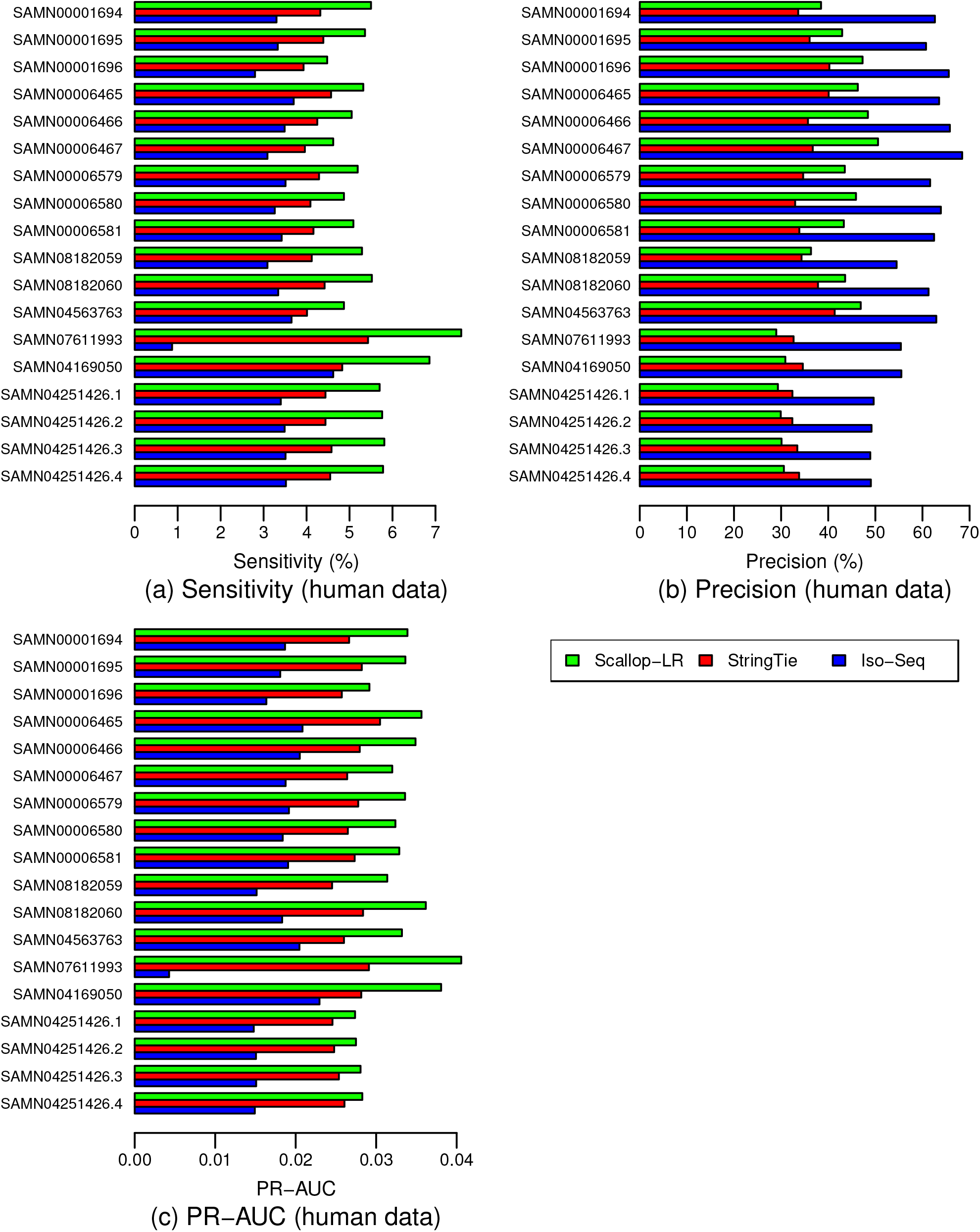
Human Data: (a) Sensitivity, (b) Precision, and (c) PR-AUC of Scallop-LR, StringTie, and Iso-Seq Analysis. Evaluations were on 18 human PacBio datasets from SRA, each corresponding to one BioSample and named by the BioSample ID (except that the last four datasets are four replicates for one BioSample). The first nine datasets were sequenced using the RS and the last nine datasets were sequenced using the RS II. Sensitivity, Precision, and PR-AUC are as described in Section 2.2

**Figure 3:**
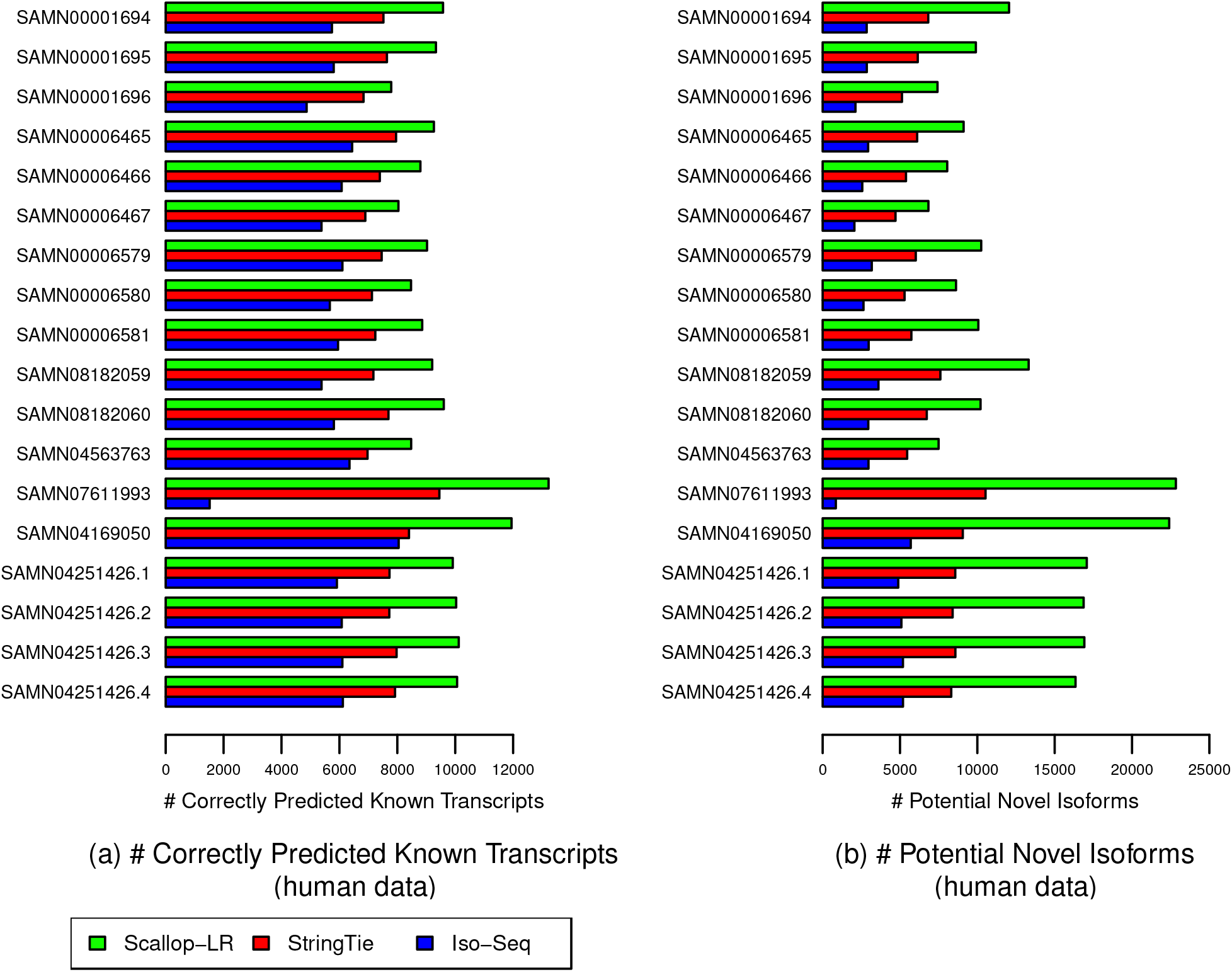
Human Data: (a) Correctly Predicted Known Transcripts, and (b) Potential Novel Isoforms of Scallop-LR, StringTie, and Iso-Seq Analysis. The same 18 human PacBio datasets as described in Figure 2 were evaluated. A Correctly Predicted Known Transcript has the exact intron-chain matching with a transcript in the reference annotation. A Potential Novel Isoform is a predicted transcript that shares at least one splice junction with a reference transcript.

In the human data, Scallop-LR also consistently assembles more known transcripts correctly than StringTie and thus consistently has higher sensitivity than StringTie. Scallop-LR finds 950– 3770 more known transcripts than StringTie, and the sensitivity of Scallop-LR is 1.14–1.42 times higher than that of StringTie (Figures 2 and 3, Tables S1 and S2). Scallop-LR’s higher sensitivity is likely due to its phasing path preservation and its transcript boundary identification in the splice graph based on the boundary information extracted from long reads.

Scallop-LR has higher precision than StringTie for the majority of the datasets. For the first 12 datasets in Figure 2 and Table S1, Scallop-LR has both higher sensitivity and higher precision than StringTie. Scallop-LR’s higher precision is partially contributed by its post-assembly clustering. However, for the last six datasets in Figure 2 and Table S1, Scallop-LR has lower precision than StringTie. The last six datasets in Figure 2 (each has 11, 12, 24, or 27 movies) are significantly larger than the first 12 datasets (each has 7 or 8 movies). Scallop-LR’s precision decreases in the six larger datasets as it assembles significantly more transcripts in total in these larger datasets (Table S2), while StringTie’s precision does not seem to change much with the size of the sample.

When two assemblers have opposite trends on sensitivity and precision on a dataset (e.g. the last six datasets in Figure 2 and Table S1), we compare their sensitivity and precision on the same footing. That is, for the assembler with a higher sensitivity, we find the precision on its PR curve by matching the sensitivity of the other assembler, and this precision is called adjusted precision. Similarly, we find the sensitivity on its PR curve by matching the precision of the other assembler, and this sensitivity is called adjusted sensitivity. The adjusted sensitivity and precision are needed only when the datasets have opposite trends on sensitivity and precision between assemblers. These adjusted values are shown inside the parentheses on Table S1. Scallop-LR’s adjusted sensitivity and adjusted precision are consistently higher than StringTie’s sensitivity and precision, indicating that Scallop-LR has consistently better performance than StringTie.

On the other hand, Iso-Seq Analysis consistently has higher precision than Scallop-LR and StringTie (Figure 2, Table S1). Iso-Seq Analysis has higher precision partially because the full-length CCS reads are run through the ICE (Iterative Clustering and Error correction) algorithm and the isoforms are also polished with Quiver to achieve higher accuracy.

Scallop-LR consistently has higher PR-AUC than Iso-Seq Analysis and StringTie, indicating better overall performance of Scallop-LR. The PR-AUC of Scallop-LR is 1.62–2.07 times higher than that of Iso-Seq Analysis, and 1.1–1.4 times higher than that of StringTie (Figure 2, Table S1).

### 3.2 Scallop-LR and StringTie find more potential novel isoforms than Iso-Seq Analysis

Scallop-LR and StringTie find more potential novel isoforms (i.e. transcripts containing at least one annotated splice junction) than Iso-Seq Analysis in the human data. Scallop-LR also consistently finds more potential novel isoforms than StringTie in the human data. Scallop-LR finds 2.53–4.23 times more potential novel isoforms than Iso-Seq Analysis, and 1.37–2.47 times more potential novel isoforms than StringTie (Figure 3, Table S2). This is likely due to the same reasons that led to the higher sensitivity of Scallop-LR. This shows the potential benefit that long-read transcript assembly could offer in discovering novel isoforms.

### 3.3 Scallop-LR finds more novel isoforms in catalog than Iso-Seq Analysis

We use SQANTI to evaluate Scallop-LR and Iso-Seq Analysis (SQANTI does not work for the transcripts assembled by StringTie). Figure 4 and Table S5 show the SQANTI evaluation results for Scallop-LR and Iso-Seq Analysis on the 18 human datasets.

**Figure 4:**
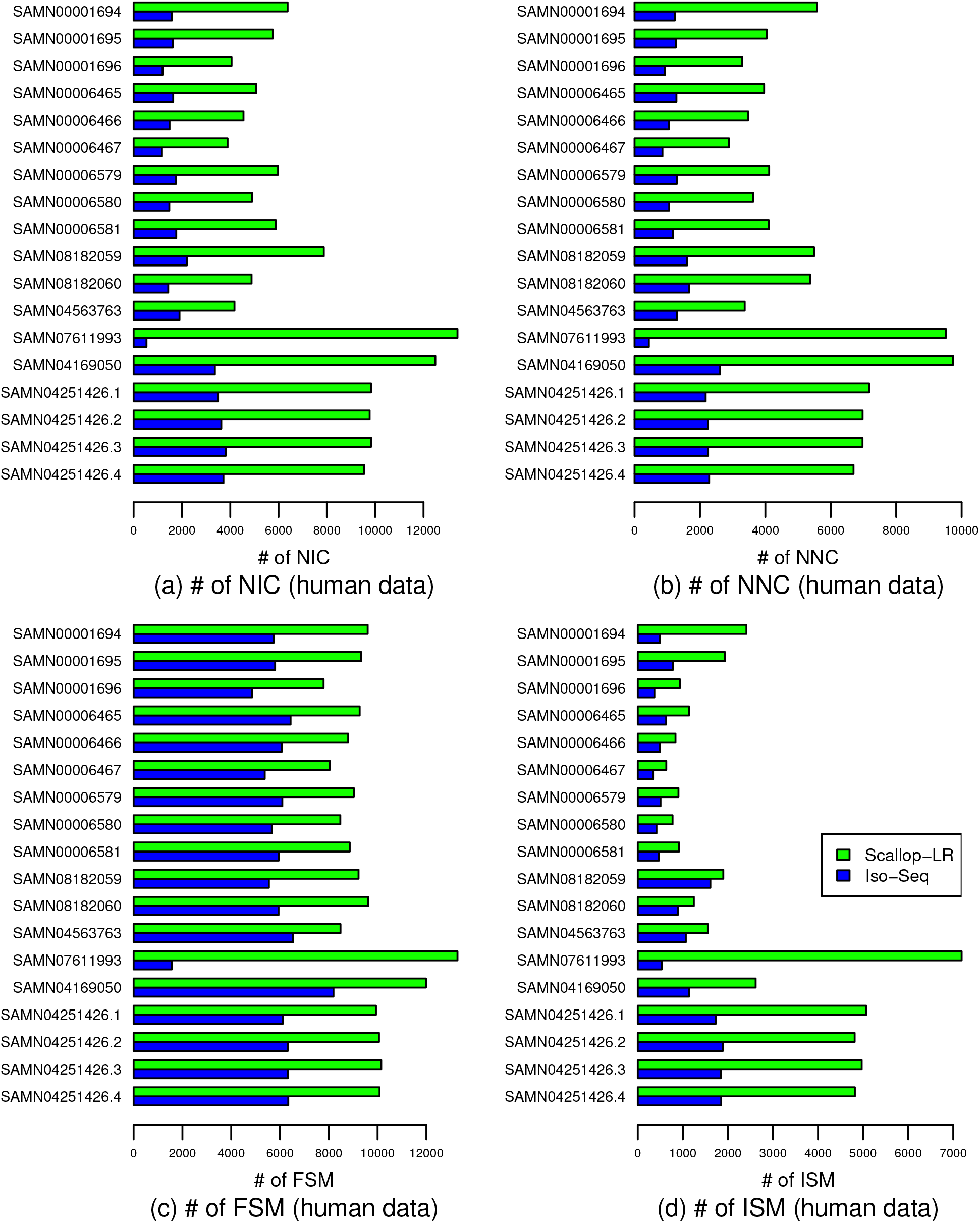
Human Data: Numbers of (a) NIC, (b) NNC, (c) FSM, and (d) ISM transcripts of Scallop-LR and Iso-Seq Analysis based on SQANTI evaluations. The same 18 human PacBio datasets as described in Figure 2 were evaluated. NIC, NNC, FSM, and ISM are as described in Section 2.2

The NIC (transcripts containing either new combinations of known splice junctions or novel splice junctions with annotated donors and acceptors) results show that Scallop-LR finds more novel isoforms in catalog than Iso-Seq Analysis consistently. Scallop-LR finds 2.2–4.02 times more NIC than Iso-Seq Analysis (Figure 4, Table S5). This is an important indication of Scallop-LR’s ability to find more new transcripts that are not yet annotated, as we conjecture that the novel isoforms in catalog may be more likely to be new transcripts than wrongly assembled transcripts since the novel splice junctions are formed from annotated donors and acceptors. This finding further supports the advantage of assembly of long reads.

The NNC (transcripts containing novel splice junctions with novel donors and/or acceptors) results indicate that Scallop-LR also finds more novel isoforms not in catalog than Iso-Seq Analysis consistently (Figure 4, Table S5). The novel isoforms not in catalog could be either new transcripts or wrongly assembled transcripts.

SQANTI’s results on novel isoforms are roughly consistent with Gffcompare’s results on novel isoforms. Comparing Table S5 with Table S2, we can see that the sums of NIC and NNC from SQANTI are similar to the numbers of potential novel isoforms reported by Gffcompare, except that for the last four datasets in Table S5, for Iso-Seq Analysis, the sums of NIC and NNC are notably larger than the corresponding numbers of potential novel isoforms in Table S2 (this may be because some NIC or NNC may not contain an annotated splice junction although they contain an annotated donor and/or acceptor).

The FSM (Full Splice Match) results from SQANTI support the trend we found from Gffcompare that Scallop-LR consistently predicts more known transcripts correctly than Iso-Seq Analysis. Comparing Table S5 with Table S2, we can see that the numbers of FSM from SQANTI are very close to the numbers of correctly predicted known transcripts from Gffcompare for these datasets.

The ISM (Incomplete Splice Match) results show that Scallop-LR also yields more partially matched transcripts than Iso-Seq Analysis (Figure 4, Table S5). The NNC and ISM results support the trend we found from Gffcompare that Iso-Seq Analysis has higher precision than Scallop-LR.

The mouse data exhibit the same trends as the human data as summarized above, which can be seen from Figure 5 and Table S6 and by comparing Table S6 with Table S4. In the mouse data, Scallop-LR finds significantly more novel isoforms in catalog (2.43–3.5 times more) than Iso-Seq Analysis consistently (Figure 5, Table S6). This further supports our finding on Scallop-LR’s ability to discover more new transcripts that are not yet annotated.

**Figure 5:**
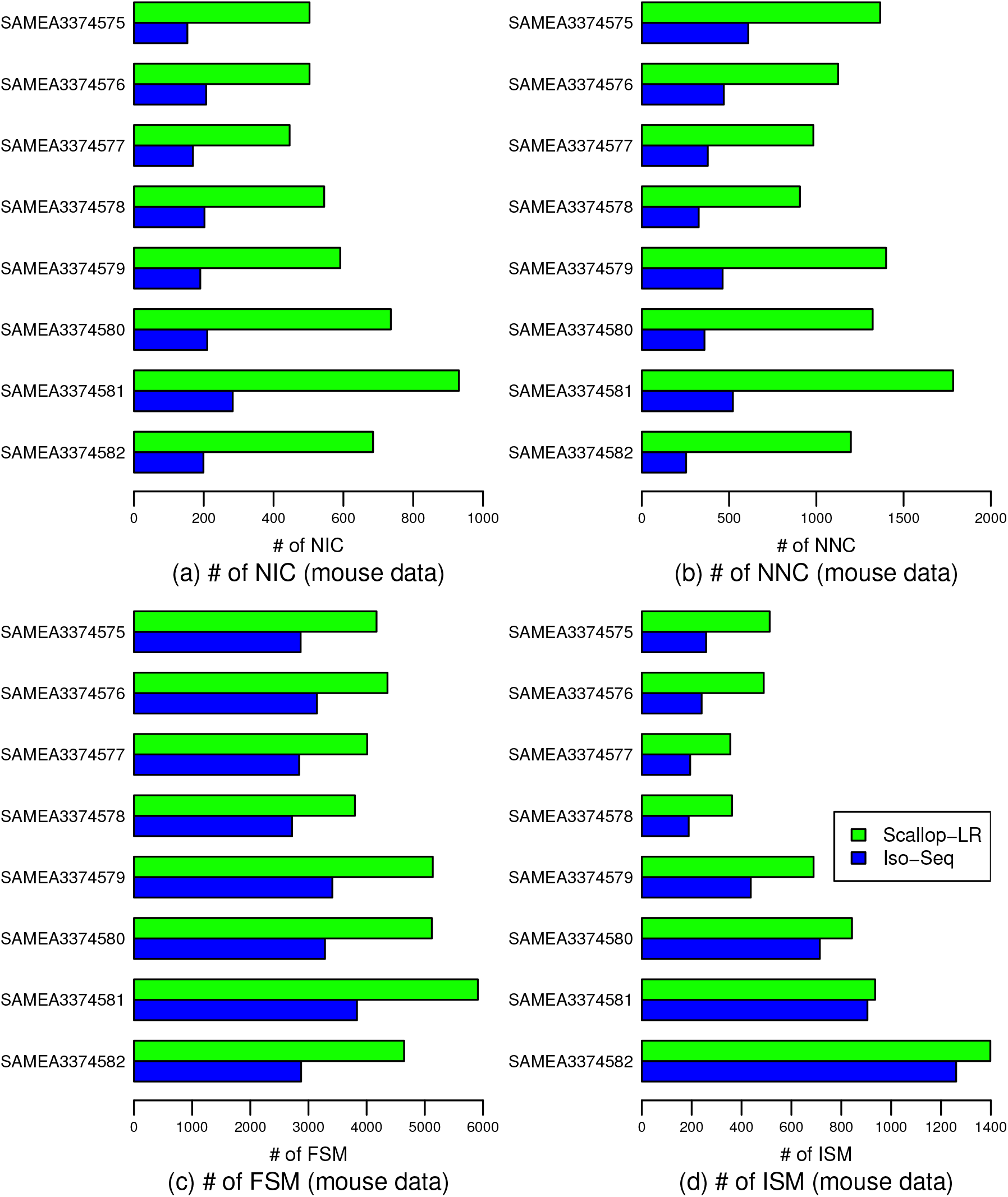
Mouse Data: Numbers of (a) NIC, (b) NNC, (c) FSM, and (d) ISM transcripts of Scallop-LR and Iso-Seq Analysis based on SQANTI evaluations. Evaluations were on eight mouse PacBio datasets from SRA, each corresponding to one BioSample and named by the BioSample ID. All eight datasets were sequenced using the RS. Metrics descriptions are the same as in Figure 4.

### 3.4 Quantification of predicted transcripts that partially match known transcripts

In rnaQUAST, “isoforms” refer to reference transcripts from the gene annotation database, and “transcripts” refer to predicted transcripts by the tools being evaluated. Here, we inherit these terminologies. Figures 6, 7, and 8 show box-whisker plots of matched transcripts in matched fraction bins, assembled isoforms in assembled fraction bins, “mean isoform assembly” and “mean fraction of transcript matched” for Scallop-LR, StringTie, and Iso-Seq Analysis on the 18 human datasets based on rnaQUAST evaluations. Full results are shown in Supplementary Tables S7.1–S7.18.

**Figure 6:**
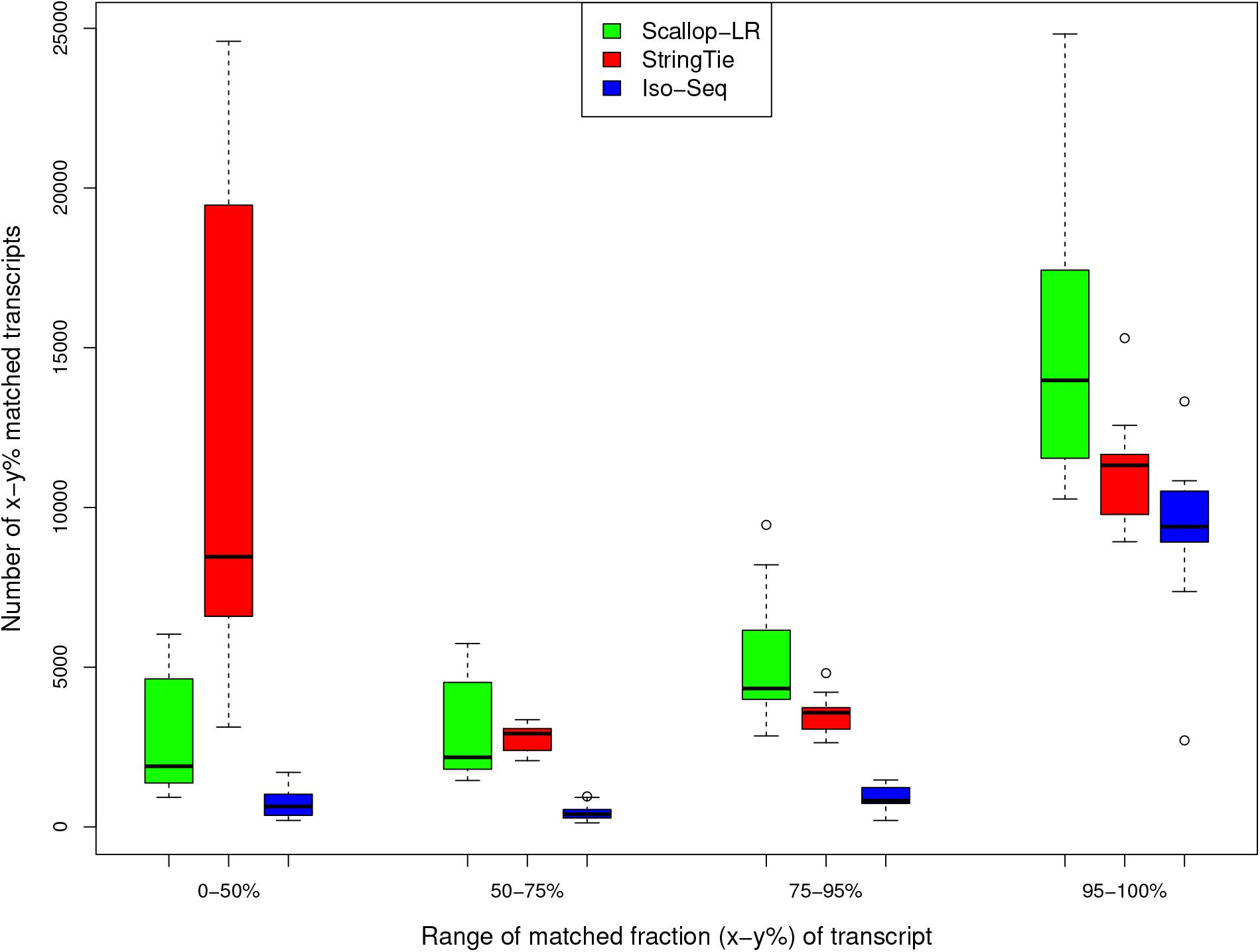
Human Data: Box-Whisker Plots of Matched Transcripts in Four Matched Fraction Bins for Scallop-LR, StringTie, and Iso-Seq Analysis, based on rnaQUAST evaluations. This is to compare numbers of x-y% matched transcripts. The same 18 human PacBio datasets as described in Figure 2 were evaluated. “Number of x-y% matched transcripts” is as described in Section 3.4. The four bins of matched fraction (x-y%) of transcript are 0–50%, 50–75%, 75–95%, and 95–100%.

**Figure 7:**
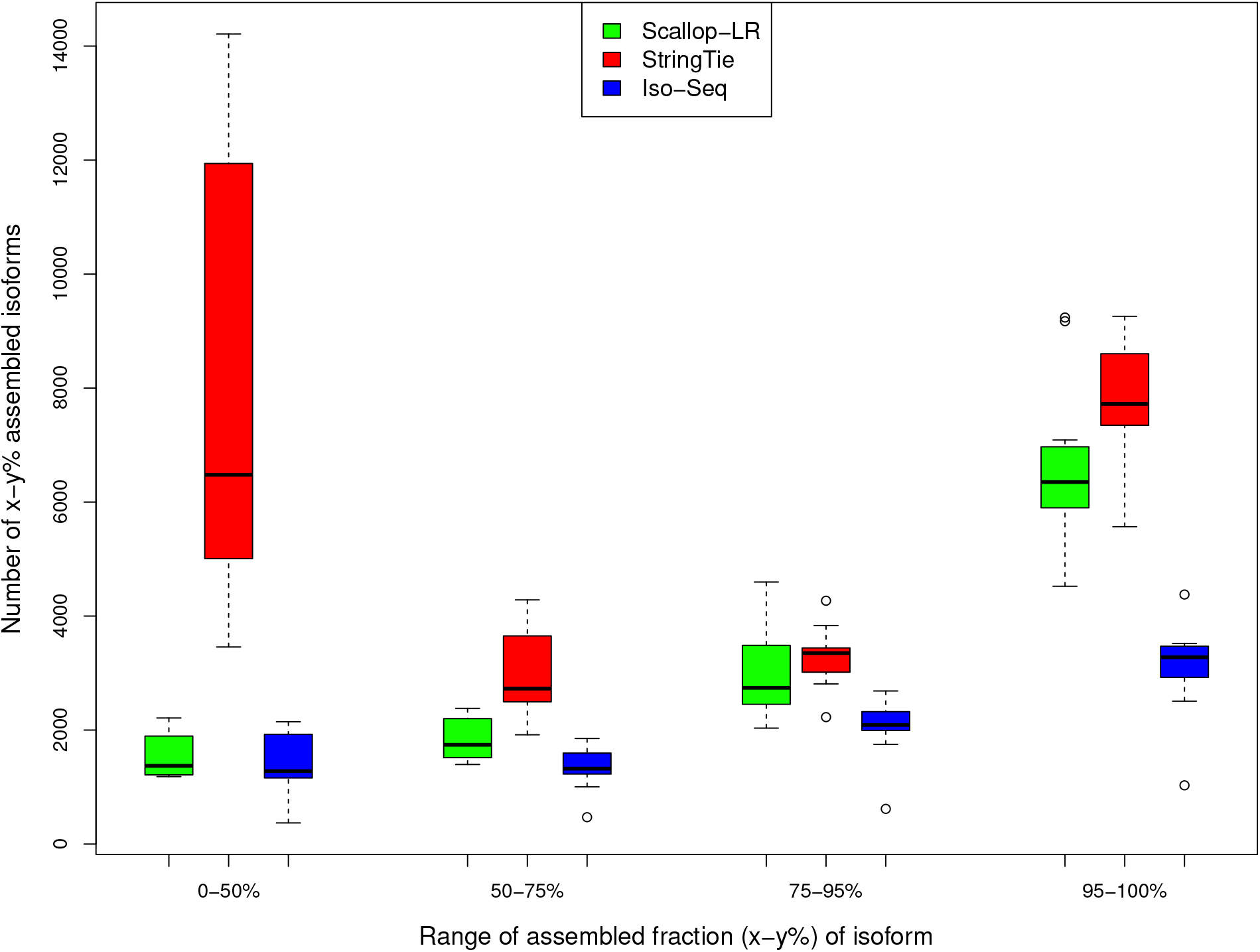
Human Data: Box-Whisker Plots of Assembled Isoforms in Four Assembled Fraction Bins for Scallop-LR, StringTie, and Iso-Seq Analysis, based on rnaQUAST evaluations. This is to compare numbers of x-y% assembled isoforms. The same 18 human PacBio datasets as described in Figure 2 were evaluated. “Number of x-y% assembled isoforms” is as described in Section 3.4. The four bins of assembled fraction (x-y%) of isoform are 0–50%, 50–75%, 75–95%, and 95–100%.

**Figure 8:**
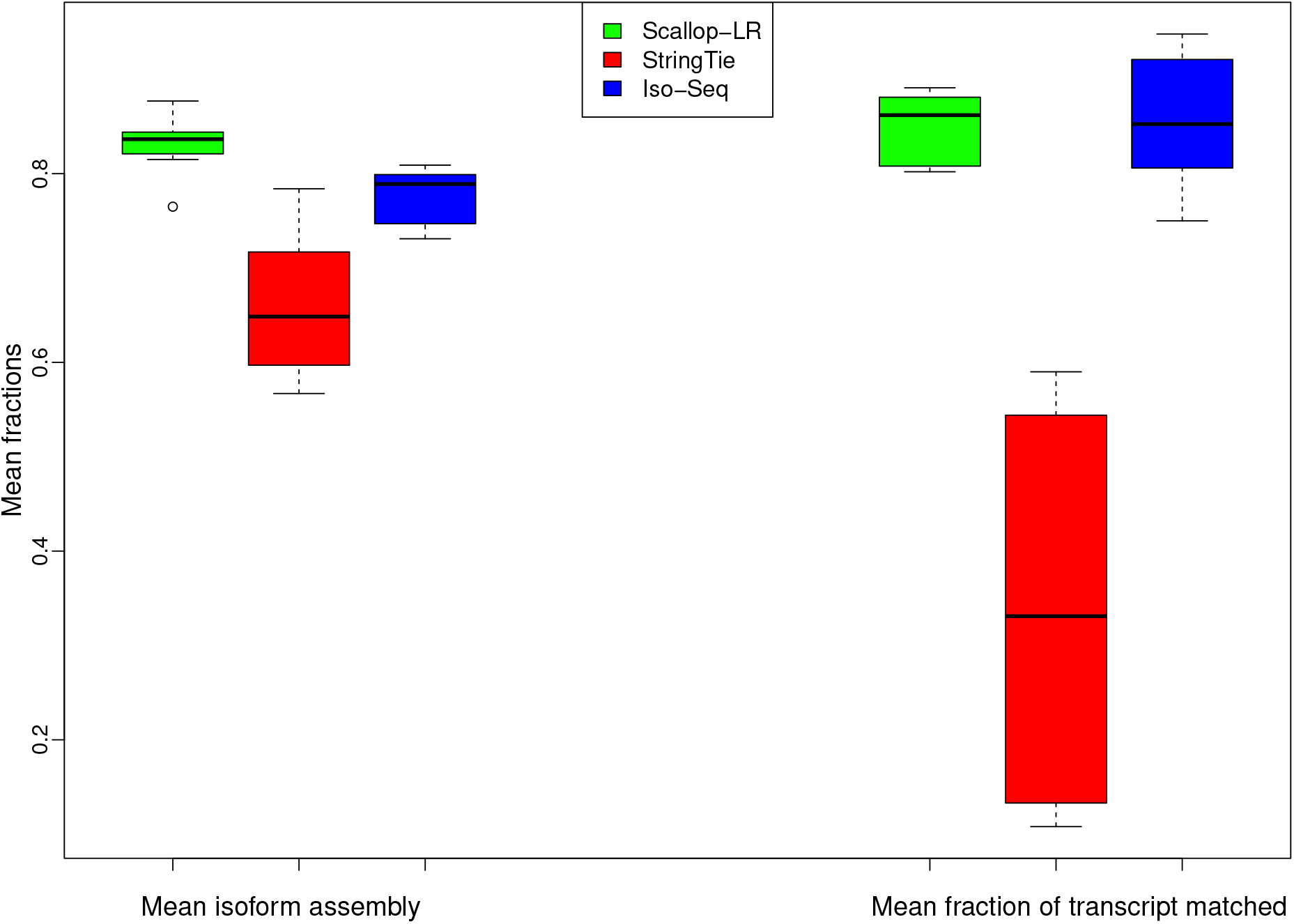
Human Data: Box-Whisker Plots of Mean Isoform Assembly and Mean Fraction of Transcript Matched for Scallop-LR, StringTie, and Iso-Seq Analysis, based on rnaQUAST evaluations. The same 18 human PacBio datasets as described in Figure 2 were evaluated. “Mean isoform assembly” and “Mean fraction of transcript matched” are as described in Section 3.4.

Scallop-LR predicts more transcripts that have a high fraction of their bases matching reference transcripts than both Iso-Seq Analysis and StringTie. The metric “x-y% matched transcripts” is the number of transcripts that have at least x% and at most y% of their bases matching an isoform from the annotation database. We report this measure in four different bins to examine how well predicted transcripts match reference transcripts. From Tables S7.1–S7.18, in the high % bins of the “x-y% matched transcripts” (75–95% and 95–100% matched), Scallop-LR predicts more x-y% matched transcripts than both Iso-Seq Analysis and StringTie (with one exception compared with StringTie). This trend is visualized in Figure 6 (75–95% and 95–100% matched bins). In the high % bins, StringTie mostly has more x-y% matched transcripts than Iso-Seq Analysis. These further support the advantage of transcript assembly on long reads.

On average, Scallop-LR transcripts match reference transcripts much better than StringTie transcripts. The metric “Mean fraction of transcript matched” is the average value of matched fractions, where the matched fraction of a transcript is computed as the number of its bases covering an isoform divided by the transcript length. This measure indicates on average how well predicted transcripts match reference transcripts. In Tables S7.1–S7.18, Scallop-LR consistently has much higher values of “Mean fraction of transcript matched” than StringTie, indicating its better assembly quality than StringTie. Scallop-LR performs slightly better than Iso-Seq Analysis on this measure. These trends are visualized in Figure 8 (right: “Mean fraction of transcript matched”).

There are more reference transcripts that have a high fraction of their bases being captured/covered by Scallop-LR transcripts than by Iso-Seq Analysis predicted transcripts. The metric “x-y% assembled isoforms” is the number of isoforms from the annotation database that have at least x% and at most y% of their bases captured by a single predicted transcript. We report this measure in four different bins to examine how well reference transcripts are captured/covered by predicted transcripts. From Tables S7.1–S7.18, in the high % bins of the “x-y% assembled isoforms” (75–95% and 95–100% assembled), Scallop-LR consistently has more x-y% assembled isoforms than Iso-Seq Analysis. However, Scallop-LR mostly (with four exceptions) has fewer x-y% assembled isoforms than StringTie in the high % bins. These trends are visualized in Figure 7 (75–95% and 95–100% assembled bins).

However, on average, reference transcripts are better captured/covered by Scallop-LR transcripts than by StringTie transcripts and Iso-Seq Analysis transcripts. The metric “Mean isoform assembly” is the average value of assembled fractions, where the assembled fraction of an isoform is computed as the largest number of its bases captured by a single predicted transcript divided by its length. This measure shows on average how well reference transcripts are captured by predicted transcripts. In Tables S7.1–S7.18, Scallop-LR consistently has higher values of “Mean isoform assembly” than both StringTie and Iso-Seq Analysis. This trend is visualized in Figure 8 (left: “Mean isoform assembly”). This trend is consistent with the higher sensitivity of Scallop-LR in the Gffcompare results.

Scallop-LR consistently has fewer unannotated, misassembled, and unaligned transcripts than StringTie (Tables S7.1–S7.18). This further indicates Scallop-LR’s better assembly quality than StringTie. Scallop-LR mostly (with three exceptions) produces fewer unannotated transcripts than Iso-Seq Analysis as well.

There are a few notable findings regarding StringTie transcripts. First, StringTie consistently has significantly more unannotated transcripts than both Scallop-LR and Iso-Seq Analysis (Tables S7.1– S7.18). Second, in Figure 6, in the 0–50% matched bin, StringTie has significantly higher numbers of transcripts than Scallop-LR and Iso-Seq Analysis. This indicates that StringTie assembled many more lower-quality transcripts than Scallop-LR and Iso-Seq Analysis, consistent with StringTie predicting many more unannotated transcripts. Lastly, in Figure 7, in the 0–50% assembled bin, StringTie has significantly higher numbers of isoforms than Scallop-LR and Iso-Seq Analysis. This indicates that, compared with Scallop-LR and Iso-Seq Analysis, there are many more isoforms from the annotation which are just marginally covered by StringTie transcripts.

The mouse data exhibit trends partially similar to those of the human data for the rnaQUAST results, and the quality of StringTie transcripts in the mouse data is somewhat improved compared to that in the human data. The detailed discussions on the rnaQUAST results for the mouse data are in the Supplementary Materials (section 3).

### 3.5 Scallop-LR and StringTie predict more known transcripts and potential novel isoforms than Iso-Seq Analysis in mouse data

From the Gffcompare evaluation for the mouse data (Figure 9, Tables S3 and S4), Scallop-LR and StringTie consistently predict more known transcripts (Scallop-LR predicts 1100–2200 more) correctly than Iso-Seq Analysis and thus consistently have higher sensitivity (Scallop-LR’s is 1.43– 1.72 times higher) than Iso-Seq Analysis. Scallop-LR and StringTie also find more potential novel isoforms (Scallop-LR finds 2.38–4.36 times more) than Iso-Seq Analysis (Table S4). Scallop-LR and StringTie consistently have higher PR-AUC than Iso-Seq Analysis (Figure 9, Table S3).

**Figure 9:**
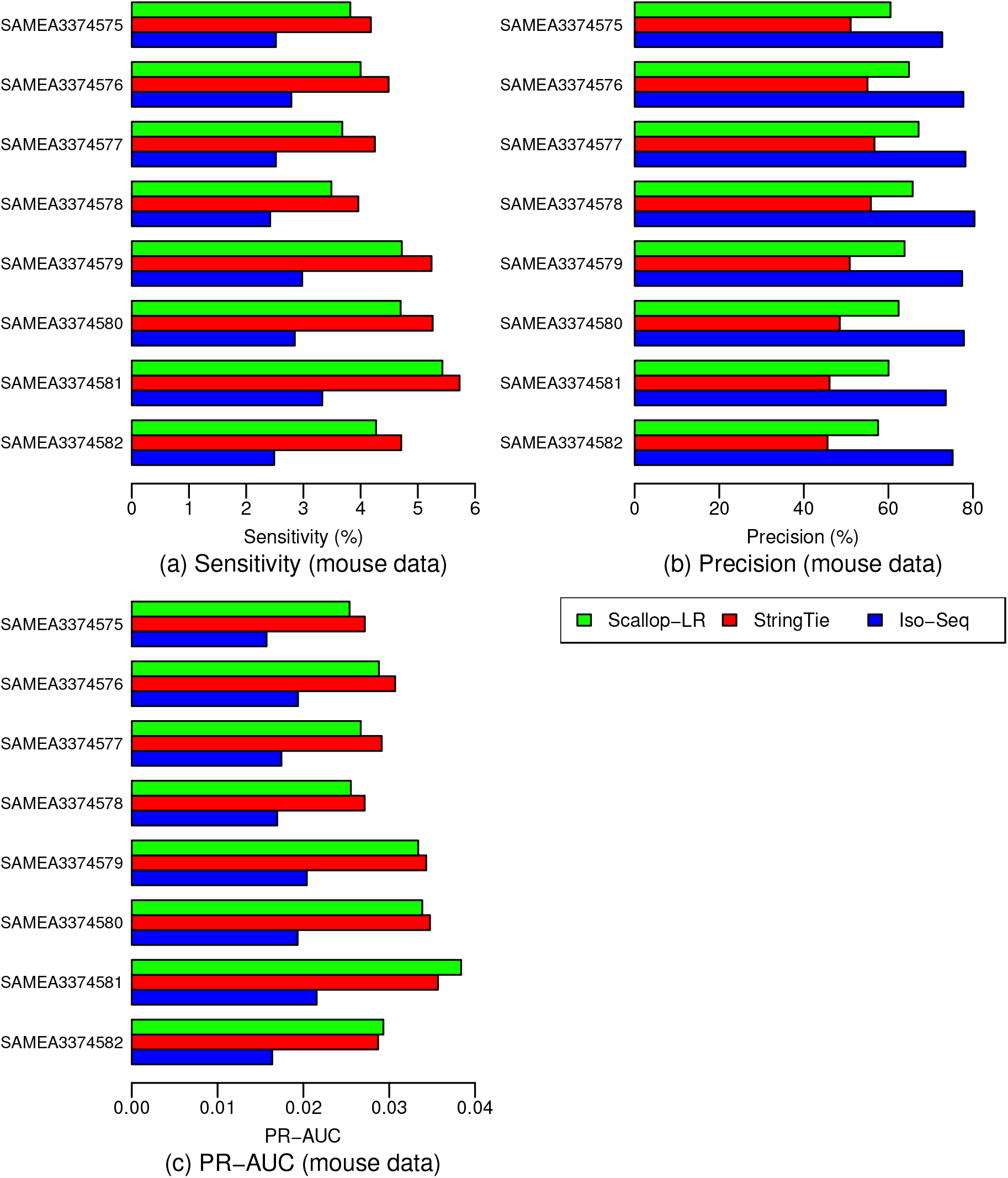
Mouse Data: (a) Sensitivity, (b) Precision, and (c) PR-AUC of Scallop-LR, StringTie, and Iso-Seq Analysis. The same eight mouse PacBio datasets as described in Figure 5 were evaluated. Metrics descriptions are the same as in Figure 2.

We also found some trends different from those in the human data. In the mouse data, Scallop-LR consistently has higher precision than StringTie, but consistently has lower sensitivity than StringTie (Figure 9, Table S3). Thus, for StringTie we computed the adjusted sensitivity by matching Scallop-LR’s precision and the adjusted precision by matching Scallop-LR’s sensitivity. These adjusted values are shown inside the parentheses on Table S3. Scallop-LR’s sensitivity and precision are consistently higher than StringTie’s adjusted sensitivity and adjusted precision, indicating that when comparing on the same footing, Scallop-LR does better on these measures than StringTie.

In the mouse data, the trend of PR-AUC between Scallop-LR and StringTie is mixed (Figure 9, Table S3). Scallop-LR also finds fewer potential novel isoforms than StringTie (Table S4).

Before this work, Scallop was never evaluated on organisms besides human, for either short reads or long reads. In fact, Scallop’s parameters were optimized by targeting the human transcriptome. The current mouse transcriptome is relatively less complex than the human transcriptome although they share many similarities. It may be possible that some of Scallop-LR’s advantages (such as preserving phasing paths) become less significant in a relatively less complex transcriptome.

## 4 Discussion

The combined evaluations using Gffcompare, SQANTI, and rnaQUAST yield consistent observations that Scallop-LR not only correctly assembles more known transcripts but also finds more possible novel isoforms than Iso-Seq Analysis, which does not do assembly. Scallop-LR finding more NIC especially shows its ability to discover new transcripts. These observations further support the idea that transcript assembly of long reads is needed and demonstrate that long-read assembly by Scallop-LR can help reveal a more complete human transcriptome using long reads.

Two factors may limit the CCS read length: the read length of the platform and the cDNA template sizes. In many cases, the primary limiting factor for CCS read lengths is the cDNA template sizes (Sharon *et al.*, 2013). When a cDNA is very long so that the continuous polymerase read is unable to get through at least two full passes of the template, the CCS read is not generated for that cDNA. Thus, the maximum possible CCS read length is limited by the read length of the platform. The read lengths of sequencing platforms have been increasing, however, there are limitations imposed by the cDNA synthesis methods.

cDNA synthesis can be incomplete with respect to the original mRNAs (Sharon *et al.*, 2013). A CCS read represents the entire cDNA molecule, however, the CCS read could correspond to a partial transcript as a result of incomplete cDNAs (Sharon *et al.*, 2013). The longer the transcripts are, the lower the fraction of CCS reads that can represent the entire splice structures of mRNAs is (Sharon *et al.*, 2013). This is likely a reason that Scallop-LR is able to find more true transcripts through assembly: a fraction of CCS reads can be partial sequences of those long transcripts, and Scallop-LR is able to assemble them together to reconstruct the original transcripts.

Iso-Seq Analysis may also sacrifice some true transcripts in order to achieve a higher quality (i.e. less affected by the sequencing errors) in final isoforms. The “polish” step in Iso-Seq Analysis keeps only the isoforms with at least two full-length reads to support them. This increases the isoform quality and gives Iso-Seq Analysis a higher precision than Scallop-LR, but may cause Iso-Seq Analysis to miss those low-abundance, long transcripts with only one full-length read.

Although StringTie was designed for assembling short reads, it also exhibits the advantage of assembly of long reads compared to Iso-Seq Analysis. StringTie finds more known transcripts and potential novel isoforms than Iso-Seq Analysis. In the rnaQUAST results, StringTie produces large numbers of unannotated transcripts (in a range of 7600–113000 for the human datasets), significantly more than those of Scallop-LR and Iso-Seq Analysis (differing by orders of magnitude). Unannotated transcripts are the transcripts that do not have a fraction matching a reference transcript in the annotation database. StringTie also outputs large numbers of single-exon transcripts, significantly more than those of Scallop-LR and Iso-Seq Analysis (differing by orders of magnitude). We found that about 70% of the unannotated transcripts from StringTie are those single-exon transcripts. StringTie produces large numbers of single-exon transcripts most likely because StringTie discards the spliced read alignments that do not have the transcript strand information. There is a fraction of read alignments by Minimap2 which have no transcript strand information, since Min-imap2 looks for the canonical splicing signal to infer the transcript strand and for some reads the transcript strands are undetermined by Minimap2. When those spliced alignments that do not have the transcript strand information are ignored by StringTie, the single-exon alignments that overlap those spliced alignments turn into single-exon transcripts by themselves, although they could have been represented by the spliced multi-exon transcripts during the assembly if those spliced alignments they overlap were not ignored. Unlike StringTie, Scallop-LR attempts both strands if a read alignment has no transcript strand information.

The sensitivity of the Iso-Seq method is limited by the factor that not all CCS reads represent full transcripts (Rhoads and Au, 2015). We demonstrate that Scallop-LR can improve this situation by identifying more true transcripts and possible novel isoforms through transcript assembly. Adding long-read-specific optimizations in Scallop-LR increases the advantage of assembly, thus providing benefit to transcriptome studies.

## Supporting information

supplementary tables, figures, and sections

## Acknowledgements

This work was supported by the T32 training grant of the US National Institutes of Health [T32 EB009403 to L.H.T.] as part of the HHMI-NIBIB Interfaces Initiative; the Gordon and Betty Moore Foundation’s Data-Driven Discovery Initiative [GBMF4554 to C.K.]; the US National Institutes of Health [R01GM122935, P41GM103712]; and The Shurl and Kay Curci Foundation.

## Footnotes

1 Pacific Biosciences. (2014). ARCHIVED: Intro to the Iso-Seq Method: Full-length transcript sequencing. June 2, 2014. https://www.pacb.com/blog/intro-to-iso-seq-method-full-leng

2 Pacific Biosciences. (2018). SMRT Tools Reference Guide v5.1.0. https://www.pacb.com/wp-content/uploads/SMRT_Tools_Reference_Guide_v510.pdf

3 The Center for Computational Biology at Johns Hopkins University. GffCompare: Program for processing GTF/GFF files. https://ccb.jhu.edu/software/stringtie/gffcompare.shtml

4 Pacific Biosciences. PacBio bioinformatics tools documentation. http://albiorix.bioenv.gu.se:8081/smrtanalysis/doc/bioinformatics-tools/GenomicConsensus/doc/HowToQuiver.html

## References

Au, K. et al. (2012). Improving PacBio long read accuracy by short read alignment. PLoS ONE, 7(10), e46679.

Au, K. et al. (2013). Characterization of the human ESC transcriptome by hybrid sequencing. PNAS, 110(50), E4821–E4830.

Bushmanova, E. et al. (2016). rnaQUAST: a quality assessment tool for de novo transcriptome assemblies. Bioinformatics, 32(14), 2210–2212.

Bushnell, B. (2014). BBMap: a fast, accurate, splice-aware aligner. 9th Annual Genomics of Energy and Environment Meeting, pages LBNL–7065E.

Cho, H. et al. (2014). High-resolution transcriptome analysis with long-read RNA sequencing. PLOS One, 9(9), e108095.

Dobin, A. et al. (2013). STAR: ultrafast universal RNA-seq aligner. Bioinformatics, 29(1), 15–21.

Hughes, J. et al. (2010). Chimpanzee and human Y chromosomes are remarkably divergent in structure and gene content. Nature, 463(7280), 536–9.

Kim, D. et al. (2013). TopHat2: accurate alignment of transcriptomes in the presence of insertions, deletions and gene fusions. Genome Biol, 14(4), R36.

Kim, D. et al. (2015). HISAT: a fast spliced aligner with low memory requirements. Nat. Methods, 12(4), 357–360.

Komor, M. et al. (2017). Identification of differentially expressed splice variants by the proteogenomic pipeline splicify. Mol Cell Proteomics, 16(10), 1850–1863.

Koren, S. et al. (2017). Canu: scalable and accurate long-read assembly via adaptive k-mer weighting and repeat separation. Genome Research, 27, 722–736.

Križanović, K. et al. (2018). Evaluation of tools for long read RNA-seq splice-aware alignment. Bioinformatics, 34(5), 748–754.

Kuosmanen, A. et al. (2016). On using longer RNA-seq reads to improve transcript prediction accuracy. 9th International Joint Conference on Biomedical Engineering Systems and Technologies, 3(BIOINFORMATICS), 272–277.

Leinonen, R. et al. (2011). The sequence read archive. Nucleic Acids Res, 39(suppL1), D19–21.

Li, H. (2017). Minimap2: fast pairwise alignment for long nucleotide sequences. arXiv, page 1708.01492v2.

O’Grady, T. et al. (2016). Global transcript structure resolution of high gene density genomes through multi-platform data integration. Nucleic Acids Res, 44(18), e145.

Pan, Q. et al. (2008). Deep surveying of alternative splicing complexity in the human transcriptome by high-throughput sequencing. Nature Genetics, 40(12), 1413–1415.

Pertea, M. et al. (2015). StringTie enables improved reconstruction of a transcriptome from RNA-seq reads. Nature Biotechnology, 33(3), 290–295.

Pertea, M. et al. (2016). Transcript-level expression analysis of RNA-seq experiments with HISAT, StringTie, and Ballgown. Nature Protocols, 11(9), 1650–1667.

Rhoads, A. and Au, K. (2015). PacBio sequencing and its applications. Genomics Proteomics Bioinformatics, 13, 278–289.

Sahlin, K. and Medvedev, P. (2019). *De novo* clustering of long-read transcriptome data using a greedy, quality-value based algorithm. RECOMB 2019, pages 227–242.

Seo, J. et al. (2016). De novo assembly and phasing of a Korean human genome. Nature, 538(7624), 243–247.

Shao, M. and Kingsford, C. (2017). Accurate assembly of transcripts through phase-preserving graph decomposition. Nature Biotechnology, 35, 1167–1169.

Sharon, D. et al. (2013). A single-molecule long-read survey of the human transcriptome. Nature Biotechnology, 31(11), 1009–1014.

Shi, L. et al. (2016). Long-read sequencing and de novo assembly of a Chinese genome. Nature Communications, 7, 12065.

Smith-Unna, R. et al. (2016). TransRate: reference-free quality assessment of de novo transcriptome assemblies. Genome Research, 26(8), 1134–1144.

Tardaguila, M. et al. (2018). SQANTI: extensive characterization of long-read transcript sequences for quality control in full-length transcriptome identification and quantification. Genome Res, 28, 396–411.

Tilgner, H. et al. (2014). Defining a personal, allele-specific, and single-molecule long-read transcriptome. PNAS, 111(27), 9869–9874.

Tseng, E. et al. (2017). Altered expression of the FMR1 splicing variants landscape in premutation carriers. Biochim Biophys Acta, 1860(11), 1117–1126.

Wang, B. et al. (2016). Unveiling the complexity of the maize transcriptome by single-molecule long-read sequencing. Nature Communications, 7, 11708.

Weirather, J. et al. (2015). Characterization of fusion genes and the significantly expressed fusion isoforms in breast cancer by hybrid sequencing. Nucleic Acids Res, 43(18), e116.

Wu, T. and Watanabe, C. (2005). GMAP: a genomic mapping and alignment program for mRNA and EST sequences. Bioinformatics, 21(9), 1859–1875.

Zimin, A. et al. (2013). The MaSuRCA genome assembler. Bioinformatics, 29(21), 2669–2677.

