## supplementary tables, figures, and sections for "Quantifying the Benefit Offered by Transcript Assembly on Single-Molecule Long Reads"

### 1. Merging multiple SRA Runs from the same BioSample into one dataset

In most of the PacBio datasets in SRA, one BioSample has multiple SRA Runs. We merge multiple SRA Runs that belong to the same BioSample into one dataset. PacBio sequencing uses the template called SMRTbell that is a closed, single-stranded circular DNA created by ligating adaptors to both ends of a target double-stranded cDNA molecule. The sequencing is based on a SMRT Cell, a chip with consumable substrates comprising arrays of zero-mode waveguide (ZMW) nanostructures, and a SMRTbell diffuses into a sequencing unit ZMW on it. The real-time observation of a SMRT Cell is called a movie, and an SRA Run usually contains one movie and sometimes contains multiple movies. One BioSample has multiple SRA Runs because the experimenters used multiple movies (i.e. multiple SMRT Cells) to increase the coverage so that those low-abundance, long isoforms can be captured in Iso-Seq Analysis, since the "polish" step in Iso-Seq Analysis keeps only the isoforms with at least two full-length reads to support them. In most cases, the experimenters also used a size selection sequencing strategy, that is, isoforms that are in different size ranges are split into separate independent SMRTbell libraries for sequencing, so that larger isoforms are not detrimentally dominated by smaller isoform molecules during the sequencing. Thus, different SRA Runs are designated for different size ranges. Therefore, we use one BioSample instead of one SRA Run to represent one dataset in our analysis, and we merge multiple SRA Runs into that dataset.

### 2. Software versions and options used in the analysis workflow

The software versions and options used in the analysis workflow are summarized in the following:

Iso-Seq Analysis: Iso-Seq2 from SMRT Link v5.1.0.
Minimap2: v2.2. Options: *-ax splice*.
StringTie: v1.3.2d. Options: *-c 1.0*.
Scallop-LR: v0.9.1. Options: *-c <ccs_read_info > --min_num_hits_in_bundle 1*.
Gffcompare: v0.9.9c. Options: *-M -N -r <reference_annotation>*.
SQANTI: v1.2. Options: *-g*.
rnaQUAST: v.1.5.1. Options: *--transcripts <multiple_assemblies> --reference <reference_genome> --gene_db <gene_database> --gmap_index <gmap_index> --labels*

*<labels> --no_plots --disable_infer_genes --disable_infer_transcripts --lower_threshold <lower_threshold> --upper_threshold <upper_threshold>*.
Transrate: v1.0.3. Options: *--assembly <assembly> --reference <reference_transcriptome>*.

### 3. Quantification of predicted transcripts that partially match known transcripts in mouse data

Figures S1, S2, and S3 show box-whisker plots of matched transcripts in matched fraction bins, assembled isoforms in assembled fraction bins, "mean isoform assembly" and "mean fraction of transcript matched" for Scallop-LR, StringTie, and Iso-Seq Analysis on the eight mouse datasets based on rnaQUAST evaluations. Full results are shown in Tables S8.1-S8.8.

In the mouse data, Scallop-LR predicts more transcripts that have a high fraction of their bases matching reference transcripts than Iso-Seq Analysis. From Tables S8.1-S8.8, in the high % bins of the “x-y% matched transcripts” (75-95% and 95-100% matched), Scallop-LR consistently has more x-y% matched transcripts than Iso-Seq Analysis. However, unlike in the human data, Scallop-LR consistently has fewer x-y% matched transcripts than StringTie in the high % bins. These trends are visualized in Figure S1 (75-95% and 95-100% matched bins).

However, on average, Scallop-LR transcripts match reference transcripts better than StringTie transcripts. In Tables S8.1-S8.8, Scallop-LR consistently has much higher values of “Mean fraction of transcript matched” than StringTie. Scallop-LR has slightly lower values than Iso-Seq Analysis though. These trends are visualized in Figure S3 (right: “Mean fraction of transcript matched”).

In the mouse data, there are more reference transcripts that have a high fraction of their bases being captured/covered by Scallop-LR transcripts than by Iso-Seq Analysis predicted transcripts. From Tables S8.1-S8.8, in the high % bins of the “x-y% assembled isoforms” (75-95% and 95-100% assembled), Scallop-LR consistently has more x-y% assembled isoforms than Iso-Seq Analysis. However, Scallop-LR consistently has fewer x-y% assembled isoforms than StringTie in the high % bins. These trends are visualized in Figure S2 (75-95% and 95-100% assembled bins).

However, on average, reference transcripts are better captured/covered by Scallop-LR transcripts than by StringTie transcripts and Iso-Seq Analysis transcripts. In Tables S8.1-S8.8, Scallop-LR consistently has higher values of “Mean isoform assembly” than both StringTie and Iso-Seq Analysis. Iso-Seq Analysis consistently has higher values than StringTie. This trend is visualized in Figure S3 (left: “Mean isoform assembly”).

The quality of StringTie transcripts in the mouse data is somewhat improved compared to that in the human data. As in the human data, StringTie consistently has significantly more unannotated transcripts than both Scallop-LR and Iso-Seq Analysis (Tables S8.1-S8.8). However, in Figure S1, unlike Figure 6, in the 0-50% matched bin StringTie no longer has a very high number of transcripts. This indicates that StringTie performs better in the mouse data than in the human data. In Figure S2, though, in the 0-50% assembled bin StringTie still has significantly higher numbers of isoforms than both Scallop-LR and Iso-Seq Analysis, similar to Figure 7.

**Table S1**: Human Data: Sensitivity, Precision, and PR-AUC of Scallop-LR, StringTie, and Iso-Seq Analysis

| Datasets | Sensitivity (%) | | | Precision (%) | | | PR-AUC | | |
|---|---|---|---|---|---|---|---|---|---|
| | Scallop-LR | StringTie | Iso-Seq | Scallop-LR | StringTie | Iso-Seq | Scallop-LR | StringTie | Iso-Seq |
| SAMN00001694 | 5.50 | 4.32 | 3.30 | 38.47 | 33.63 | 62.64 | 0.03389 | 0.02664 | 0.01868 |
| SAMN00001695 | 5.36 | 4.39 | 3.33 | 42.96 | 36.05 | 60.76 | 0.03363 | 0.02821 | 0.01808 |
| SAMN00001696 | 4.48 | 3.93 | 2.80 | 47.29 | 40.24 | 65.59 | 0.02916 | 0.02573 | 0.01636 |
| SAMN00006465 | 5.32 | 4.57 | 3.70 | 46.28 | 40.10 | 63.54 | 0.03563 | 0.03048 | 0.02085 |
| SAMN00006466 | 5.05 | 4.25 | 3.49 | 48.42 | 35.70 | 65.82 | 0.03489 | 0.02796 | 0.02051 |
| SAMN00006467 | 4.62 | 3.96 | 3.09 | 50.57 | 36.71 | 68.43 | 0.03200 | 0.02640 | 0.01875 |
| SAMN00006579 | 5.19 | 4.29 | 3.51 | 43.52 | 34.68 | 61.63 | 0.03359 | 0.02777 | 0.01916 |
| SAMN00006580 | 4.87 | 4.09 | 3.26 | 45.89 | 32.98 | 63.91 | 0.03239 | 0.02648 | 0.01839 |
| SAMN00006581 | 5.09 | 4.16 | 3.42 | 43.31 | 33.87 | 62.47 | 0.03287 | 0.02733 | 0.01906 |
| SAMN08182059 | 5.29 | 4.12 | 3.09 | 36.34 | 34.34 | 54.52 | 0.03138 | 0.02452 | 0.01515 |
| SAMN08182060 | 5.52 | 4.42 | 3.34 | 43.59 | 37.82 | 61.30 | 0.03617 | 0.02837 | 0.01832 |
| SAMN04563763 | 4.87 | 4.01 | 3.65 | 46.90 | 41.37 | 62.94 | 0.03320 | 0.02600 | 0.02047 |
| SAMN07611993 | 7.60 (7.26) | 5.43 | 0.87 | 28.97 (46.47) | 32.65 | 55.42 | 0.04057 | 0.02910 | 0.00427 |
| SAMN04169050 | 6.86 (6.61) | 4.83 | 4.62 | 30.90 (51.74) | 34.61 | 55.52 | 0.03807 | 0.02815 | 0.02296 |
| SAMN04251426.1 | 5.70 (5.46) | 4.44 | 3.40 | 29.32 (40.72) | 32.40 | 49.64 | 0.02738 | 0.02457 | 0.01479 |
| SAMN04251426.2 | 5.76 (5.56) | 4.44 | 3.49 | 29.92 (41.18) | 32.39 | 49.19 | 0.02749 | 0.02478 | 0.01507 |
| SAMN04251426.3 | 5.81 (5.52) | 4.58 | 3.51 | 30.09 (40.37) | 33.44 | 48.93 | 0.02806 | 0.02537 | 0.01509 |
| SAMN04251426.4 | 5.78 (5.47) | 4.55 | 3.52 | 30.59 (40.91) | 33.82 | 49.05 | 0.02828 | 0.02606 | 0.01492 |

The above table compares the Gffcompare evaluation results for Scallop-LR, StringTie, and Iso-Seq Analysis on human data. 18 human PacBio datasets were extracted from SRA, each corresponding to one BioSample and named by the BioSample ID (except that the last four datasets are four replicates for one BioSample). Multiple SRA Runs that belong to each BioSample were merged into a large dataset to perform the analyses. The first nine datasets were sequenced using the RS instrument and the last nine datasets were sequenced using the RS II instrument. Sensitivity is the ratio of the number of correctly predicted known transcripts over the total number of known transcripts, and precision is the ratio of the number of correctly predicted known transcripts over the total number of predicted transcripts. PR-AUC was calculated from the precision-recall curves we generated. The values within the parentheses are the adjusted sensitivity and adjusted precision. The adjusted sensitivity for Scallop-LR was calculated by matching the precision of StringTie, and the adjusted precision for Scallop-LR was calculated by matching the sensitivity of StringTie. The adjusted sensitivity and precision were only calculated for the last six datasets, since the last six datasets have opposite trends on sensitivity and precision comparing Scallop-LR and StringTie.

**Table S2**: Human Data: Correctly Predicted Known Transcripts, Total Multi-Exon Transcripts, and Potential Novel Isoforms of Scallop-LR, StringTie, and Iso-Seq Analysis

| Datasets | # Potential Novel Isoforms | | | # Total Multi-Exon Transcripts | | | # Correctly Predicted Known Transcripts | | |
|---|---|---|---|---|---|---|---|---|---|
| | Scallop-LR | StringTie | Iso-Seq | Scallop-LR | StringTie | Iso-Seq | Scallop-LR | StringTie | Iso-Seq |
| SAMN00001694 | 12050 | 6827 | 2847 | 24903 | 22370 | 9166 | 9580 | 7522 | 5742 |
| SAMN00001695 | 9905 | 6149 | 2856 | 21731 | 21201 | 9554 | 9336 | 7642 | 5805 |
| SAMN00001696 | 7425 | 5129 | 2122 | 16476 | 16983 | 7423 | 7791 | 6834 | 4869 |
| SAMN00006465 | 9112 | 6111 | 2941 | 20019 | 19847 | 10136 | 9264 | 7958 | 6440 |
| SAMN00006466 | 8054 | 5387 | 2561 | 18171 | 20715 | 9236 | 8798 | 7396 | 6079 |
| SAMN00006467 | 6838 | 4710 | 2054 | 15900 | 18783 | 7865 | 8040 | 6896 | 5382 |
| SAMN00006579 | 10250 | 6020 | 3175 | 20742 | 21508 | 9906 | 9027 | 7460 | 6105 |
| SAMN00006580 | 8623 | 5295 | 2640 | 18467 | 21607 | 8870 | 8474 | 7125 | 5669 |
| SAMN00006581 | 10064 | 5736 | 2974 | 20458 | 21376 | 9531 | 8861 | 7241 | 5954 |
| SAMN08182059 | 13318 | 7610 | 3609 | 25332 | 20893 | 9876 | 9206 | 7175 | 5384 |
| SAMN08182060 | 10207 | 6732 | 2951 | 22038 | 20348 | 9478 | 9606 | 7696 | 5810 |
| SAMN04563763 | 7496 | 5464 | 2965 | 18078 | 16857 | 10087 | 8478 | 6973 | 6349 |
| SAMN07611993 | 22834 | 10532 | 852 | 45657 | 28953 | 2741 | 13226 | 9453 | 1519 |
| SAMN04169050 | 22403 | 9059 | 5696 | 38657 | 24280 | 14493 | 11946 | 8403 | 8047 |
| SAMN04251426.1 | 17074 | 8572 | 4887 | 33824 | 23863 | 11913 | 9916 | 7731 | 5914 |
| SAMN04251426.2 | 16871 | 8403 | 5090 | 33534 | 23849 | 12363 | 10034 | 7724 | 6081 |
| SAMN04251426.3 | 16916 | 8580 | 5191 | 33637 | 23850 | 12472 | 10122 | 7976 | 6103 |
| SAMN04251426.4 | 16347 | 8314 | 5194 | 32908 | 23423 | 12476 | 10065 | 7922 | 6119 |

The above table compares additional Gffcompare evaluation results for Scallop-LR, StringTie, and Iso-Seq Analysis on human data. The same 18 human PacBio datasets as described in Table S1 were evaluated. A Correctly Predicted Known Transcript is a transcript that has the exact intron-chain matching with a transcript in the reference annotation. A Potential Novel Isoform is a predicted transcript that shares at least one splice junction with a reference transcript. The # Total Multi-Exon Transcripts is the total number of predicted multi-exon transcripts.

**Table S3**: Mouse Data: Sensitivity, Precision, and PR-AUC of Scallop-LR, StringTie, and Iso-Seq Analysis

| Datasets | Sensitivity (%) | | | Precision (%) | | | PR-AUC | | |
|---|---|---|---|---|---|---|---|---|---|
| | Scallop-LR | StringTie | Iso-Seq | Scallop-LR | StringTie | Iso-Seq | Scallop-LR | StringTie | Iso-Seq |
| SAMEA3374575 | 3.82 | 4.18 (3.70) | 2.52 | 60.50 | 51.08 (57.55) | 72.74 | 0.02538 | 0.02716 | 0.01571 |
| SAMEA3374576 | 4.00 | 4.49 (3.87) | 2.79 | 64.86 | 55.06 (62.48) | 77.74 | 0.02880 | 0.03069 | 0.01937 |
| SAMEA3374577 | 3.68 | 4.25 (3.61) | 2.52 | 67.14 | 56.71 (65.44) | 78.23 | 0.02668 | 0.02913 | 0.01744 |
| SAMEA3374578 | 3.49 | 3.96 (3.38) | 2.42 | 65.74 | 55.86 (64.21) | 80.37 | 0.02553 | 0.02714 | 0.01695 |
| SAMEA3374579 | 4.72 | 5.24 (4.48) | 2.98 | 63.83 | 50.86 (58.13) | 77.44 | 0.03338 | 0.03431 | 0.02040 |
| SAMEA3374580 | 4.70 | 5.26 (4.52) | 2.85 | 62.42 | 48.50 (58.28) | 77.87 | 0.03385 | 0.03475 | 0.01933 |
| SAMEA3374581 | 5.43 | 5.73 (4.93) | 3.33 | 60.04 | 46.09 (50.63) | 73.58 | 0.03838 | 0.03568 | 0.02155 |
| SAMEA3374582 | 4.27 | 4.71 (4.05) | 2.49 | 57.57 | 45.62 (53.16) | 75.21 | 0.02932 | 0.02870 | 0.01637 |

This table compares the Gffcompare evaluation results for Scallop-LR, StringTie, and Iso-Seq Analysis on mouse data. Eight mouse PacBio datasets were extracted from SRA, each corresponding to one BioSample and named by the BioSample ID. Multiple SRA Runs that belong to each BioSample were merged into a large dataset to perform the analyses. All eight datasets were sequenced using the RS instrument. Sensitivity, precision, and PR-AUC are as described in Table S1. The values within the parentheses are the adjusted sensitivity and adjusted precision. The adjusted sensitivity for StringTie was calculated by matching the precision of Scallop-LR, and the adjusted precision for StringTie was calculated by matching the sensitivity of Scallop-LR. The adjusted sensitivity and precision were calculated for all eight datasets, since all eight datasets have opposite trends on sensitivity and precision comparing Scallop-LR and StringTie.

**Table S4**: Mouse Data: Correctly Predicted Known Transcripts, Total Multi-Exon Transcripts, and Potential Novel Isoforms of Scallop-LR, StringTie, and Iso-Seq Analysis

| Datasets | # Potential Novel Isoforms | | | # Total Multi-Exon Transcripts | | | # Correctly Predicted Known Transcripts | | |
|---|---|---|---|---|---|---|---|---|---|
| | Scallop-LR | StringTie | Iso-Seq | Scallop-LR | StringTie | Iso-Seq | Scallop-LR | StringTie | Iso-Seq |
| SAMEA3374575 | 1973 | 2476 | 829 | 6879 | 8913 | 3768 | 4162 | 4553 | 2741 |
| SAMEA3374576 | 1671 | 2206 | 699 | 6707 | 8870 | 3903 | 4350 | 4884 | 3034 |
| SAMEA3374577 | 1468 | 1973 | 596 | 5962 | 8155 | 3505 | 4003 | 4625 | 2742 |
| SAMEA3374578 | 1434 | 1958 | 520 | 5774 | 7723 | 3280 | 3796 | 4314 | 2636 |
| SAMEA3374579 | 2003 | 2973 | 663 | 8048 | 11221 | 4193 | 5137 | 5707 | 3247 |
| SAMEA3374580 | 2020 | 3237 | 543 | 8204 | 11801 | 3981 | 5121 | 5724 | 3100 |
| SAMEA3374581 | 2714 | 3806 | 832 | 9842 | 13531 | 4920 | 5909 | 6237 | 3620 |
| SAMEA3374582 | 1803 | 2934 | 414 | 8069 | 11233 | 3606 | 4645 | 5125 | 2712 |

This table compares additional Gffcompare evaluation results for Scallop-LR, StringTie, and Iso-Seq Analysis on mouse data. The same eight mouse PacBio datasets as described in Table S3 were evaluated. The Correctly Predicted Known Transcript, Potential Novel Isoform, and # Total Multi-Exon Transcripts are as described in Table S2.

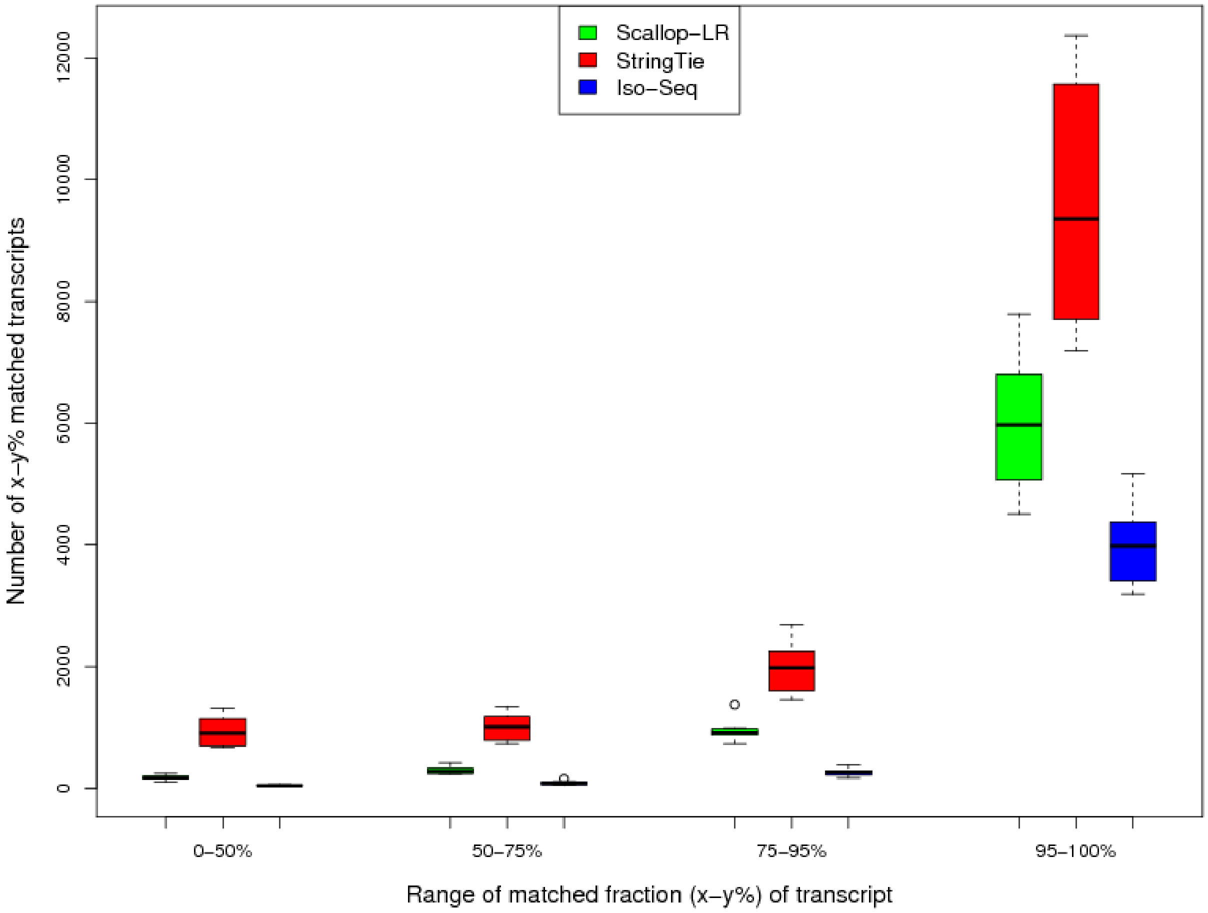


**Figure S1**: Mouse Data: Box-Whisker Plots of Matched Transcripts in Four Matched Fraction Bins for Scallop-LR, StringTie, and Iso-Seq Analysis, based on rnaQUAST evaluations. This figure is to compare numbers of x-y% matched transcripts. The same eight mouse PacBio datasets as described in Table S3 were evaluated via rnaQUAST. Figure axis descriptions are the same as in Figure 6.

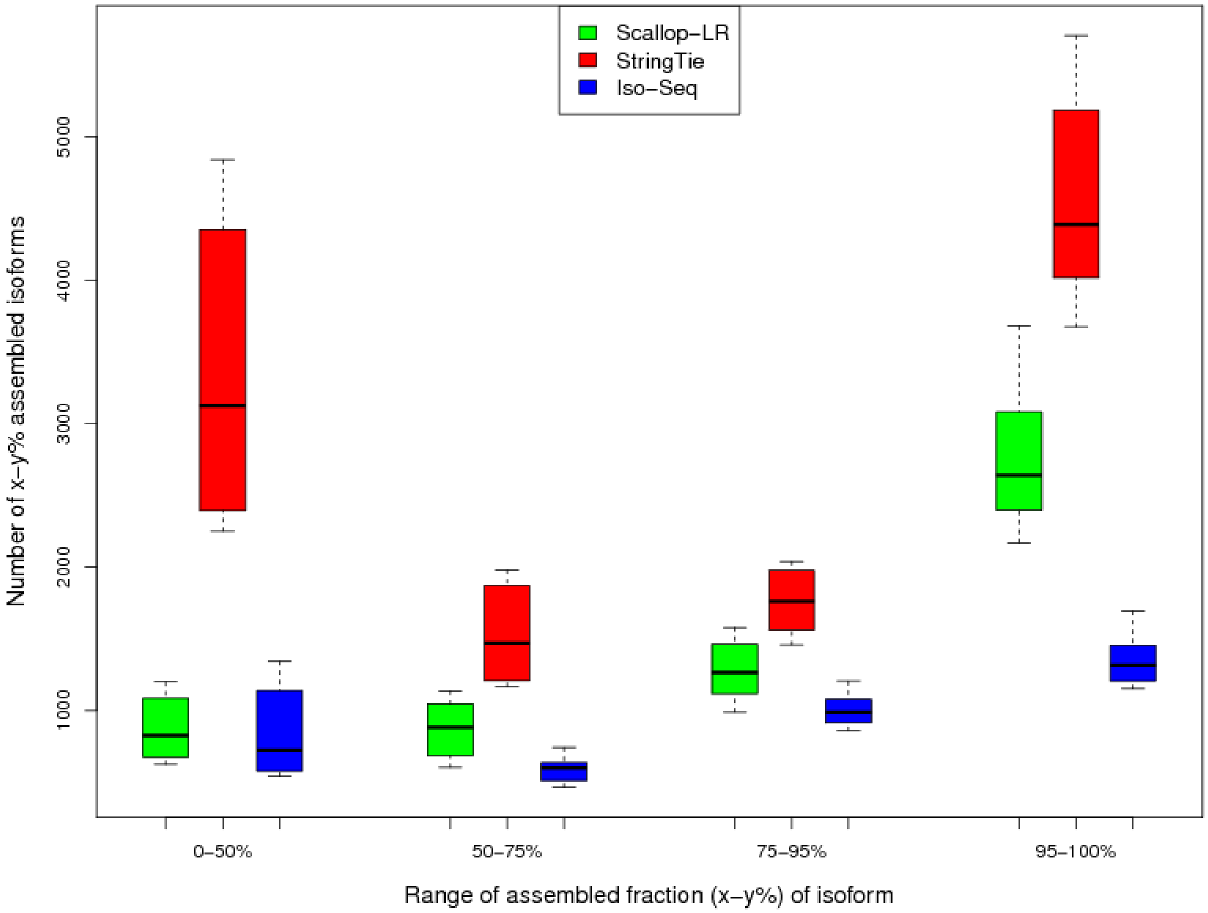


**Figure S2**: Mouse Data: Box-Whisker Plots of Assembled Isoforms in Four Assembled Fraction Bins for Scallop-LR, StringTie, and Iso-Seq Analysis, based on rnaQUAST evaluations. This figure is to compare numbers of x-y% assembled isoforms. The same eight mouse PacBio datasets as described in Table S3 were evaluated via rnaQUAST. Figure axis descriptions are the same as in Figure 7.

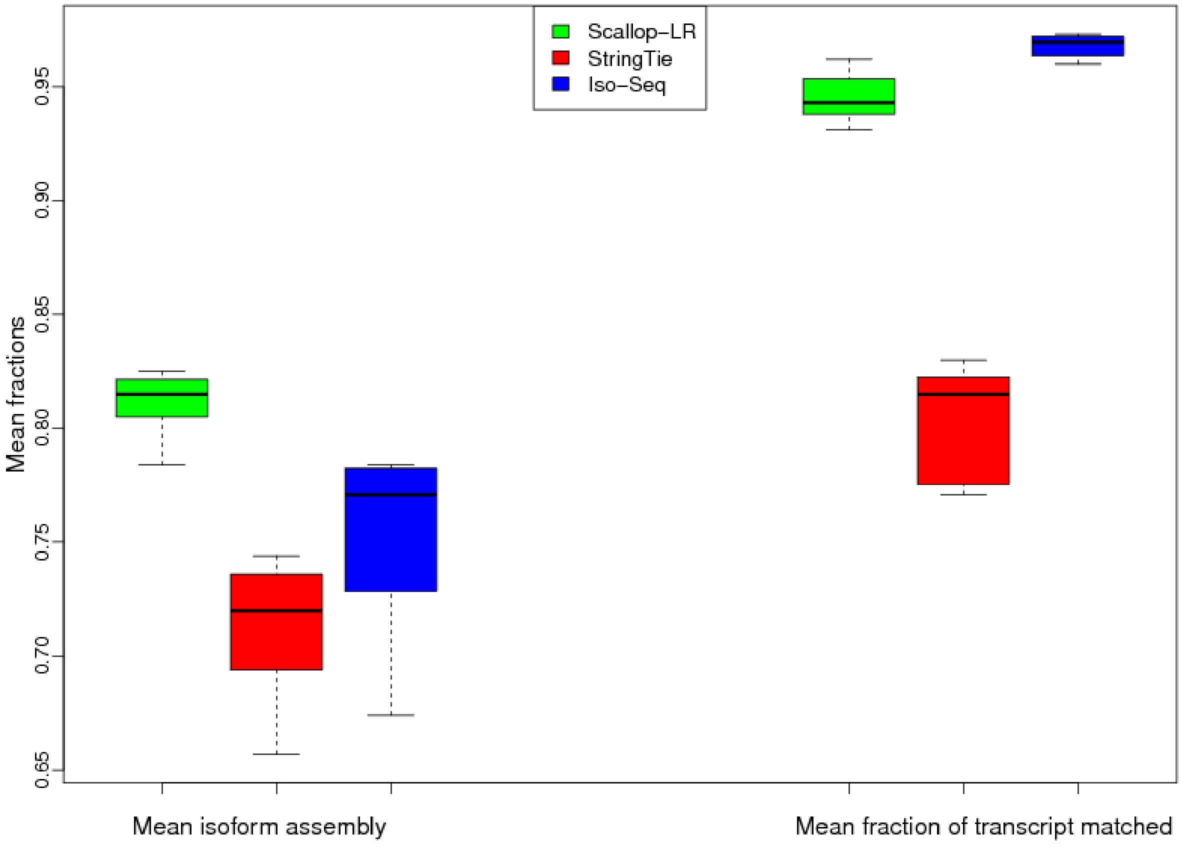


**Figure S3**: Mouse Data: Box-Whisker Plots of Mean Isoform Assembly and Mean Fraction of Transcript Matched for Scallop-LR, StringTie, and Iso-Seq Analysis, based on rnaQUAST evaluations. The same eight mouse PacBio datasets as described in Table S3 were evaluated via rnaQUAST. Figure axis descriptions are the same as in Figure 8.

**Table S5**: Human Data: Numbers of FSM, NIC, NNC, and ISM transcripts of Scallop-LR and Iso-Seq Analysis based on SQANTI Evaluations

| Datasets | # of FSM | | # of NIC | | # of NNC | | # of ISM | |
|---|---|---|---|---|---|---|---|---|
| | Scallop-LR | Iso-Seq | Scallop-LR | Iso-Seq | Scallop-LR | Iso-Seq | Scallop-LR | Iso-Seq |
| SAMN00001694 | 9594 | 5736 | 6375 | 1586 | 5579 | 1232 | 2411 | 487 |
| SAMN00001695 | 9329 | 5800 | 5765 | 1624 | 4045 | 1263 | 1934 | 776 |
| SAMN00001696 | 7787 | 4861 | 4054 | 1198 | 3294 | 930 | 934 | 369 |
| SAMN00006465 | 9266 | 6434 | 5080 | 1638 | 3962 | 1281 | 1141 | 630 |
| SAMN00006466 | 8796 | 6071 | 4548 | 1488 | 3479 | 1057 | 837 | 493 |
| SAMN00006467 | 8039 | 5375 | 3890 | 1173 | 2890 | 854 | 633 | 340 |
| SAMN00006579 | 9026 | 6096 | 5979 | 1759 | 4115 | 1297 | 903 | 505 |
| SAMN00006580 | 8472 | 5665 | 4903 | 1478 | 3628 | 1060 | 772 | 415 |
| SAMN00006581 | 8859 | 5947 | 5889 | 1765 | 4104 | 1176 | 919 | 470 |
| SAMN08182059 | 9220 | 5545 | 7874 | 2206 | 5488 | 1611 | 1898 | 1612 |
| SAMN08182060 | 9616 | 5941 | 4882 | 1433 | 5375 | 1674 | 1244 | 888 |
| SAMN04563763 | 8480 | 6535 | 4174 | 1899 | 3368 | 1297 | 1556 | 1066 |
| SAMN07611993 | 13275 | 1564 | 13406 | 533 | 9518 | 438 | 7186 | 531 |
| SAMN04169050 | 11984 | 8193 | 12490 | 3364 | 9737 | 2616 | 2616 | 1141 |
| SAMN04251426.1 | 9934 | 6114 | 9831 | 3497 | 7174 | 2175 | 5070 | 1729 |
| SAMN04251426.2 | 10058 | 6317 | 9771 | 3630 | 6972 | 2248 | 4813 | 1886 |
| SAMN04251426.3 | 10150 | 6327 | 9830 | 3814 | 6970 | 2249 | 4969 | 1840 |
| SAMN04251426.4 | 10081 | 6340 | 9545 | 3719 | 6695 | 2282 | 4818 | 1851 |

The above table compares the SQANTI evaluation results for Scallop-LR and Iso-Seq Analysis on human data. The same 18 human PacBio datasets as described in Table S1 were evaluated. FSM (Full Splice Match): the predicted transcript that matches a reference transcript at all splice junctions. ISM (Incomplete Splice Match): the predicted transcript that matches consecutive, but not all, splice junctions of a reference transcript. NIC (Novel in Catalog): the predicted transcript that contains new combinations of already annotated splice junctions or novel splice junctions formed from already annotated donors and acceptors. NNC (Novel Not in Catalog): the predicted transcript that contains novel splice junctions formed from novel donors or/and novel acceptors.

**Table S6**: Mouse Data: Numbers of FSM, NIC, NNC, and ISM transcripts of Scallop-LR and Iso-Seq Analysis based on SQANTI Evaluations

| Datasets | # of FSM | | # of NIC | | # of NNC | | # of ISM | |
|---|---|---|---|---|---|---|---|---|
| | Scallop-LR | Iso-Seq | Scallop-LR | Iso-Seq | Scallop-LR | Iso-Seq | Scallop-LR | Iso-Seq |
| SAMEA3374575 | 4170 | 2865 | 503 | 153 | 1366 | 610 | 513 | 258 |
| SAMEA3374576 | 4358 | 3145 | 503 | 207 | 1125 | 470 | 489 | 240 |
| SAMEA3374577 | 4009 | 2840 | 446 | 169 | 983 | 378 | 355 | 194 |
| SAMEA3374578 | 3799 | 2718 | 545 | 202 | 906 | 326 | 362 | 188 |
| SAMEA3374579 | 5137 | 3410 | 591 | 190 | 1400 | 463 | 689 | 437 |
| SAMEA3374580 | 5121 | 3284 | 736 | 210 | 1323 | 359 | 843 | 714 |
| SAMEA3374581 | 5912 | 3834 | 931 | 283 | 1784 | 522 | 936 | 905 |
| SAMEA3374582 | 4644 | 2874 | 685 | 199 | 1198 | 254 | 1398 | 1261 |

This table compares the SQANTI evaluation results for Scallop-LR and Iso-Seq Analysis on mouse data. The same eight mouse PacBio datasets as described in Table S3 were evaluated. Metrics descriptions are the same as in Table S5.

**Table S7.1**: rnaQUAST evaluation results for human dataset SAMN08182059, comparing Scallop-LR, StringTie, and Iso-Seq Analysis

| Metrics | Scallop-LR | StringTie | Iso-Seq |
|---|---|---|---|
| # Transcripts | 25364 | 32230 | 11467 |
| Aligned | 25363 | 32224 | 11467 |
| Uniquely aligned | 25361 | 32028 | 11444 |
| Unaligned | 1 | 6 | 0 |
| Misassemblies | 2 | 4 | 4 |
| 0-50% assembled isoforms | 1412 | 4156 | 1681 |
| 50-75% assembled isoforms | 1827 | 2332 | 1358 |
| 75-95% assembled isoforms | 2994 | 3012 | 1994 |
| 95-100% assembled isoforms | 6052 | 6691 | 2507 |
| Mean isoform assembly | 0.834 | 0.729 | 0.74 |
| 0-50% matched transcripts | 1917 | 4466 | 267 |
| 50-75% matched transcripts | 2057 | 2394 | 231 |
| 75-95% matched transcripts | 4612 | 3659 | 734 |
| 95-100% matched transcripts | 16595 | 12570 | 9971 |
| Unannotated | 180 | 9118 | 260 |
| Mean fraction of transcript matched | 0.883 | 0.558 | 0.939 |

This table compares the rnaQUAST evaluation results for Scallop-LR, StringTie, and Iso-Seq Analysis on a human dataset. "Isoforms" refer to reference transcripts from the gene annotation database, and "transcripts" refer to predicted transcripts. "# Transcripts" is the total number of predicted transcripts (including single-exon transcripts). "Aligned" is the number of transcripts which have at least one significant alignment to the reference genome. "Uniquely aligned" is the number of transcripts which have a single significant alignment. "Unaligned" is the number of transcripts without any significant alignments. "Misassemblies" are the transcripts that have discordant best-scored alignments (i.e. partial alignments that are mapped to different strands, different chromosomes, in reverse order, or too far away). "Unannotated" is the number of transcripts that do not cover any isoform from the annotation database. "x-y% assembled isoforms" is the number of isoforms from the annotation database that have at least x% and at most y% captured by a single predicted transcript. "x-y% matched transcripts" is the number of transcripts that have at least x% and at most y% matching an isoform from the annotation database. "Mean isoform assembly" is the average value of assembled fractions, where the assembled fraction of an isoform is computed as the largest number of its bases captured by a single predicted transcript divided by its length. "Mean fraction of transcript matched" is the average value of matched fractions, where the matched fraction of a transcript is computed as the number of its bases covering an isoform divided by the transcript length.

**Table S7.2**: rnaQUAST evaluation results for human dataset SAMN08182060, comparing Scallop-LR, StringTie, and Iso-Seq Analysis

| Metrics | Scallop-LR | StringTie | Iso-Seq |
|---|---|---|---|
| # Transcripts | 22065 | 29664 | 10233 |
| Aligned | 22065 | 29660 | 10233 |
| Uniquely aligned | 22057 | 29480 | 10219 |
| Unaligned | 0 | 4 | 0 |
| Misassemblies | 3 | 4 | 2 |
| 0-50% assembled isoforms | 1205 | 3659 | 1154 |
| 50-75% assembled isoforms | 1547 | 1994 | 1236 |
| 75-95% assembled isoforms | 2759 | 2810 | 2035 |
| 95-100% assembled isoforms | 6760 | 7420 | 3064 |
| Mean isoform assembly | 0.855 | 0.757 | 0.794 |
| 0-50% matched transcripts | 1677 | 4263 | 266 |
| 50-75% matched transcripts | 1806 | 2147 | 186 |
| 75-95% matched transcripts | 4035 | 3588 | 652 |
| 95-100% matched transcripts | 14376 | 11914 | 9004 |
| Unannotated | 165 | 7726 | 123 |
| Mean fraction of transcript matched | 0.881 | 0.577 | 0.948 |

This table compares the rnaQUAST evaluation results for Scallop-LR, StringTie, and Iso-Seq Analysis on a human dataset. Metrics descriptions are the same as in Table S7.1.

**Table S7.3**: rnaQUAST evaluation results for human dataset SAMN04563763, comparing Scallop-LR, StringTie, and Iso-Seq Analysis

| Metrics | Scallop-LR | StringTie | Iso-Seq |
|---|---|---|---|
| # Transcripts | 18097 | 29003 | 11103 |
| Aligned | 18096 | 29002 | 11103 |
| Uniquely aligned | 18093 | 28788 | 11088 |
| Unaligned | 1 | 1 | 0 |
| Misassemblies | 2 | 3 | 0 |
| 0-50% assembled isoforms | 2212 | 5762 | 1926 |
| 50-75% assembled isoforms | 1792 | 2496 | 1411 |
| 75-95% assembled isoforms | 2034 | 2227 | 1954 |
| 95-100% assembled isoforms | 4523 | 5567 | 2669 |
| Mean isoform assembly | 0.765 | 0.657 | 0.731 |
| 0-50% matched transcripts | 927 | 3126 | 362 |
| 50-75% matched transcripts | 1657 | 2337 | 327 |
| 75-95% matched transcripts | 4318 | 3769 | 1166 |
| 95-100% matched transcripts | 11053 | 11733 | 9105 |
| Unannotated | 138 | 8017 | 143 |
| Mean fraction of transcript matched | 0.89 | 0.59 | 0.934 |

This table compares the rnaQUAST evaluation results for Scallop-LR, StringTie, and Iso-Seq Analysis on a human dataset. Metrics descriptions are the same as in Table S7.1.

**Table S7.4**: rnaQUAST evaluation results for human dataset SAMN07611993, comparing Scallop-LR, StringTie, and Iso-Seq Analysis

| Metrics | Scallop-LR | StringTie | Iso-Seq |
|---|---|---|---|
| # Transcripts | 45773 | 42040 | 3382 |
| Aligned | 45769 | 42028 | 3382 |
| Uniquely aligned | 45763 | 41813 | 3378 |
| Unaligned | 4 | 12 | 0 |
| Misassemblies | 5 | 9 | 3 |
| 0-50% assembled isoforms | 1572 | 3951 | 371 |
| 50-75% assembled isoforms | 2380 | 2458 | 471 |
| 75-95% assembled isoforms | 4597 | 4269 | 617 |
| 95-100% assembled isoforms | 9173 | 9143 | 1030 |
| Mean isoform assembly | 0.855 | 0.775 | 0.795 |
| 0-50% matched transcripts | 5156 | 6578 | 204 |
| 50-75% matched transcripts | 5739 | 3356 | 126 |
| 75-95% matched transcripts | 9454 | 4814 | 203 |
| 95-100% matched transcripts | 24821 | 15300 | 2709 |
| Unannotated | 594 | 11959 | 137 |
| Mean fraction of transcript matched | 0.832 | 0.544 | 0.883 |

This table compares the rnaQUAST evaluation results for Scallop-LR, StringTie, and Iso-Seq Analysis on a human dataset. Metrics descriptions are the same as in Table S7.1.

**Table S7.5**: rnaQUAST evaluation results for human dataset SAMN04169050, comparing Scallop-LR, StringTie, and Iso-Seq Analysis

| Metrics | Scallop-LR | StringTie | Iso-Seq |
|---|---|---|---|
| # Transcripts | 38737 | 33225 | 15752 |
| Aligned | 38734 | 33217 | 15752 |
| Uniquely aligned | 38727 | 33099 | 15750 |
| Unaligned | 3 | 8 | 0 |
| Misassemblies | 6 | 7 | 0 |
| 0-50% assembled isoforms | 1203 | 3458 | 1387 |
| 50-75% assembled isoforms | 1691 | 1917 | 1596 |
| 75-95% assembled isoforms | 3377 | 3015 | 2686 |
| 95-100% assembled isoforms | 9238 | 8800 | 4378 |
| Mean isoform assembly | 0.877 | 0.784 | 0.809 |
| 0-50% matched transcripts | 6034 | 6593 | 580 |
| 50-75% matched transcripts | 5428 | 3086 | 506 |
| 75-95% matched transcripts | 8205 | 4216 | 1232 |
| 95-100% matched transcripts | 18839 | 11650 | 13320 |
| Unannotated | 221 | 7662 | 114 |
| Mean fraction of transcript matched | 0.802 | 0.558 | 0.939 |

This table compares the rnaQUAST evaluation results for Scallop-LR, StringTie, and Iso-Seq Analysis on a human dataset. Metrics descriptions are the same as in Table S7.1.

**Table S7.6**: rnaQUAST evaluation results for human dataset SAMN04251426.1, comparing Scallop-LR, StringTie, and Iso-Seq Analysis

| Metrics | Scallop-LR | StringTie | Iso-Seq |
|---|---|---|---|
| # Transcripts | 33883 | 41154 | 15342 |
| Aligned | 33882 | 41147 | 15342 |
| Uniquely aligned | 33881 | 40883 | 15322 |
| Unaligned | 1 | 7 | 0 |
| Misassemblies | 1 | 19 | 10 |
| 0-50% assembled isoforms | 1972 | 5067 | 1999 |
| 50-75% assembled isoforms | 2334 | 2666 | 1812 |
| 75-95% assembled isoforms | 3649 | 3429 | 2323 |
| 95-100% assembled isoforms | 6903 | 7347 | 3350 |
| Mean isoform assembly | 0.815 | 0.714 | 0.747 |
| 0-50% matched transcripts | 4722 | 7576 | 1554 |
| 50-75% matched transcripts | 4542 | 3059 | 865 |
| 75-95% matched transcripts | 6157 | 3734 | 1385 |
| 95-100% matched transcripts | 17879 | 11661 | 10297 |
| Unannotated | 581 | 15084 | 1228 |
| Mean fraction of transcript matched | 0.807 | 0.442 | 0.806 |

This table compares the rnaQUAST evaluation results for Scallop-LR, StringTie, and Iso-Seq Analysis on a human dataset. Metrics descriptions are the same as in Table S7.1.

**Table S7.7**: rnaQUAST evaluation results for human dataset SAMN04251426.2, comparing Scallop-LR, StringTie, and Iso-Seq Analysis

| Metrics | Scallop-LR | StringTie | Iso-Seq |
|---|---|---|---|
| # Transcripts | 33588 | 41008 | 16119 |
| Aligned | 33587 | 41004 | 16119 |
| Uniquely aligned | 33578 | 40814 | 16104 |
| Unaligned | 1 | 4 | 0 |
| Misassemblies | 6 | 11 | 7 |
| 0-50% assembled isoforms | 1903 | 5087 | 2051 |
| 50-75% assembled isoforms | 2201 | 2609 | 1827 |
| 75-95% assembled isoforms | 3602 | 3276 | 2418 |
| 95-100% assembled isoforms | 7051 | 7377 | 3477 |
| Mean isoform assembly | 0.821 | 0.713 | 0.748 |
| 0-50% matched transcripts | 4834 | 7801 | 1615 |
| 50-75% matched transcripts | 4525 | 3048 | 925 |
| 75-95% matched transcripts | 6174 | 3653 | 1393 |
| 95-100% matched transcripts | 17429 | 11337 | 10787 |
| Unannotated | 616 | 15143 | 1389 |
| Mean fraction of transcript matched | 0.802 | 0.435 | 0.8 |

This table compares the rnaQUAST evaluation results for Scallop-LR, StringTie, and Iso-Seq Analysis on a human dataset. Metrics descriptions are the same as in Table S7.1.

**Table S7.8**: rnaQUAST evaluation results for human dataset SAMN04251426.3, comparing Scallop-LR, StringTie, and Iso-Seq Analysis

| Metrics | Scallop-LR | StringTie | Iso-Seq |
|---|---|---|---|
| # Transcripts | 33699 | 41038 | 16328 |
| Aligned | 33698 | 41036 | 16328 |
| Uniquely aligned | 33691 | 40815 | 16306 |
| Unaligned | 1 | 2 | 0 |
| Misassemblies | 6 | 16 | 4 |
| 0-50% assembled isoforms | 1894 | 5007 | 2146 |
| 50-75% assembled isoforms | 2236 | 2616 | 1834 |
| 75-95% assembled isoforms | 3486 | 3416 | 2417 |
| 95-100% assembled isoforms | 7089 | 7384 | 3471 |
| Mean isoform assembly | 0.821 | 0.717 | 0.744 |
| 0-50% matched transcripts | 4595 | 7693 | 1707 |
| 50-75% matched transcripts | 4538 | 3008 | 955 |
| 75-95% matched transcripts | 6329 | 3757 | 1469 |
| 95-100% matched transcripts | 17653 | 11587 | 10747 |
| Unannotated | 576 | 14965 | 1441 |
| Mean fraction of transcript matched | 0.808 | 0.441 | 0.794 |

This table compares the rnaQUAST evaluation results for Scallop-LR, StringTie, and Iso-Seq Analysis on a human dataset. Metrics descriptions are the same as in Table S7.1.

**Table S7.9**: rnaQUAST evaluation results for human dataset SAMN04251426.4, comparing Scallop-LR, StringTie, and Iso-Seq Analysis

| Metrics | Scallop-LR | StringTie | Iso-Seq |
|---|---|---|---|
| # Transcripts | 32952 | 40754 | 16179 |
| Aligned | 32951 | 40749 | 16179 |
| Uniquely aligned | 32943 | 40528 | 16157 |
| Unaligned | 1 | 5 | 0 |
| Misassemblies | 7 | 11 | 7 |
| 0-50% assembled isoforms | 1940 | 5087 | 2109 |
| 50-75% assembled isoforms | 2246 | 2729 | 1852 |
| 75-95% assembled isoforms | 3563 | 3361 | 2430 |
| 95-100% assembled isoforms | 6970 | 7220 | 3414 |
| Mean isoform assembly | 0.818 | 0.711 | 0.743 |
| 0-50% matched transcripts | 4634 | 7569 | 1647 |
| 50-75% matched transcripts | 4322 | 3079 | 910 |
| 75-95% matched transcripts | 6116 | 3565 | 1440 |
| 95-100% matched transcripts | 17260 | 11484 | 10837 |
| Unannotated | 611 | 15037 | 1334 |
| Mean fraction of transcript matched | 0.806 | 0.438 | 0.803 |

This table compares the rnaQUAST evaluation results for Scallop-LR, StringTie, and Iso-Seq Analysis on a human dataset. Metrics descriptions are the same as in Table S7.1.

**Table S7.10**: rnaQUAST evaluation results for human dataset SAMN00001694, comparing Scallop-LR, StringTie, and Iso-Seq Analysis

| Metrics | Scallop-LR | StringTie | Iso-Seq |
|---|---|---|---|
| # Transcripts | 24956 | 81557 | 10813 |
| Aligned | 24956 | 81554 | 10813 |
| Uniquely aligned | 24948 | 81117 | 10802 |
| Unaligned | 0 | 3 | 0 |
| Misassemblies | 7 | 9 | 0 |
| 0-50% assembled isoforms | 1547 | 8224 | 1070 |
| 50-75% assembled isoforms | 2016 | 3162 | 1263 |
| 75-95% assembled isoforms | 2967 | 3329 | 2025 |
| 95-100% assembled isoforms | 6470 | 7697 | 3107 |
| Mean isoform assembly | 0.83 | 0.64 | 0.802 |
| 0-50% matched transcripts | 2015 | 11463 | 466 |
| 50-75% matched transcripts | 2582 | 2822 | 398 |
| 75-95% matched transcripts | 5034 | 3726 | 762 |
| 95-100% matched transcripts | 15137 | 11144 | 8916 |
| Unannotated | 180 | 52379 | 270 |
| Mean fraction of transcript matched | 0.868 | 0.219 | 0.915 |

This table compares the rnaQUAST evaluation results for Scallop-LR, StringTie, and Iso-Seq Analysis on a human dataset. Metrics descriptions are the same as in Table S7.1.

**Table S7.11**: rnaQUAST evaluation results for human dataset SAMN00001695, comparing Scallop-LR, StringTie, and Iso-Seq Analysis

| Metrics | Scallop-LR | StringTie | Iso-Seq |
|---|---|---|---|
| # Transcripts | 21741 | 107273 | 12057 |
| Aligned | 21741 | 107269 | 12057 |
| Uniquely aligned | 21736 | 106575 | 12039 |
| Unaligned | 0 | 4 | 0 |
| Misassemblies | 3 | 15 | 1 |
| 0-50% assembled isoforms | 1469 | 10422 | 1269 |
| 50-75% assembled isoforms | 1808 | 3535 | 1356 |
| 75-95% assembled isoforms | 2723 | 3400 | 2179 |
| 95-100% assembled isoforms | 6144 | 8079 | 3138 |
| Mean isoform assembly | 0.832 | 0.607 | 0.786 |
| 0-50% matched transcripts | 1498 | 15153 | 606 |
| 50-75% matched transcripts | 2146 | 2924 | 367 |
| 75-95% matched transcripts | 4350 | 3529 | 829 |
| 95-100% matched transcripts | 13582 | 10679 | 9484 |
| Unannotated | 161 | 74942 | 770 |
| Mean fraction of transcript matched | 0.879 | 0.165 | 0.872 |

This table compares the rnaQUAST evaluation results for Scallop-LR, StringTie, and Iso-Seq Analysis on a human dataset. Metrics descriptions are the same as in Table S7.1.

**Table S7.12**: rnaQUAST evaluation results for human dataset SAMN00001696, comparing Scallop-LR, StringTie, and Iso-Seq Analysis

| Metrics | Scallop-LR | StringTie | Iso-Seq |
|---|---|---|---|
| # Transcripts | 16487 | 65425 | 8855 |
| Aligned | 16487 | 65418 | 8855 |
| Uniquely aligned | 16482 | 65094 | 8847 |
| Unaligned | 0 | 7 | 0 |
| Misassemblies | 3 | 4 | 1 |
| 0-50% assembled isoforms | 1183 | 7195 | 859 |
| 50-75% assembled isoforms | 1482 | 2726 | 1004 |
| 75-95% assembled isoforms | 2364 | 2904 | 1749 |
| 95-100% assembled isoforms | 5115 | 6335 | 2604 |
| Mean isoform assembly | 0.834 | 0.633 | 0.807 |
| 0-50% matched transcripts | 1066 | 9110 | 350 |
| 50-75% matched transcripts | 1454 | 2074 | 240 |
| 75-95% matched transcripts | 3011 | 2636 | 573 |
| 95-100% matched transcripts | 10875 | 9927 | 7369 |
| Unannotated | 77 | 41656 | 321 |
| Mean fraction of transcript matched | 0.891 | 0.227 | 0.911 |

This table compares the rnaQUAST evaluation results for Scallop-LR, StringTie, and Iso-Seq Analysis on a human dataset. Metrics descriptions are the same as in Table S7.1.

**Table S7.13**: rnaQUAST evaluation results for human dataset SAMN00006465, comparing Scallop-LR, StringTie, and Iso-Seq Analysis

| Metrics | Scallop-LR | StringTie | Iso-Seq |
|---|---|---|---|
| # Transcripts | 20038 | 82621 | 12557 |
| Aligned | 20038 | 82617 | 12557 |
| Uniquely aligned | 20033 | 82068 | 12546 |
| Unaligned | 0 | 4 | 0 |
| Misassemblies | 4 | 11 | 1 |
| 0-50% assembled isoforms | 1333 | 8262 | 1180 |
| 50-75% assembled isoforms | 1630 | 2921 | 1348 |
| 75-95% assembled isoforms | 2517 | 3083 | 2176 |
| 95-100% assembled isoforms | 6315 | 7746 | 3520 |
| Mean isoform assembly | 0.842 | 0.636 | 0.804 |
| 0-50% matched transcripts | 1375 | 11130 | 426 |
| 50-75% matched transcripts | 1856 | 2375 | 397 |
| 75-95% matched transcripts | 3993 | 3258 | 854 |
| 95-100% matched transcripts | 12688 | 11297 | 10512 |
| Unannotated | 122 | 54521 | 367 |
| Mean fraction of transcript matched | 0.882 | 0.207 | 0.921 |

This table compares the rnaQUAST evaluation results for Scallop-LR, StringTie, and Iso-Seq Analysis on a human dataset. Metrics descriptions are the same as in Table S7.1.

**Table S7.14**: rnaQUAST evaluation results for human dataset SAMN00006466, comparing Scallop-LR, StringTie, and Iso-Seq Analysis

| Metrics | Scallop-LR | StringTie | Iso-Seq |
|---|---|---|---|
| # Transcripts | 18192 | 140280 | 12335 |
| Aligned | 18190 | 140274 | 12335 |
| Uniquely aligned | 18187 | 139647 | 12307 |
| Unaligned | 2 | 6 | 0 |
| Misassemblies | 1 | 18 | 1 |
| 0-50% assembled isoforms | 1315 | 13372 | 1414 |
| 50-75% assembled isoforms | 1502 | 3942 | 1229 |
| 75-95% assembled isoforms | 2481 | 3485 | 2072 |
| 95-100% assembled isoforms | 5899 | 8561 | 3280 |
| Mean isoform assembly | 0.84 | 0.574 | 0.782 |
| 0-50% matched transcripts | 1300 | 20842 | 701 |
| 50-75% matched transcripts | 1649 | 3109 | 391 |
| 75-95% matched transcripts | 3547 | 3061 | 749 |
| 95-100% matched transcripts | 11541 | 9780 | 9300 |
| Unannotated | 151 | 103450 | 1193 |
| Mean fraction of transcript matched | 0.879 | 0.122 | 0.833 |

This table compares the rnaQUAST evaluation results for Scallop-LR, StringTie, and Iso-Seq Analysis on a human dataset. Metrics descriptions are the same as in Table S7.1.

**Table S7.15**: rnaQUAST evaluation results for human dataset SAMN00006467, comparing Scallop-LR, StringTie, and Iso-Seq Analysis

| Metrics | Scallop-LR | StringTie | Iso-Seq |
|---|---|---|---|
| # Transcripts | 15914 | 134501 | 10487 |
| Aligned | 15914 | 134492 | 10487 |
| Uniquely aligned | 15912 | 133856 | 10458 |
| Unaligned | 0 | 9 | 0 |
| Misassemblies | 2 | 17 | 0 |
| 0-50% assembled isoforms | 1199 | 13214 | 1160 |
| 50-75% assembled isoforms | 1396 | 3910 | 1081 |
| 75-95% assembled isoforms | 2322 | 3441 | 1846 |
| 95-100% assembled isoforms | 5296 | 8069 | 2925 |
| Mean isoform assembly | 0.839 | 0.567 | 0.79 |
| 0-50% matched transcripts | 1202 | 20694 | 674 |
| 50-75% matched transcripts | 1463 | 2767 | 281 |
| 75-95% matched transcripts | 2848 | 2744 | 574 |
| 95-100% matched transcripts | 10264 | 8927 | 7765 |
| Unannotated | 135 | 99314 | 1192 |
| Mean fraction of transcript matched | 0.878 | 0.117 | 0.813 |

This table compares the rnaQUAST evaluation results for Scallop-LR, StringTie, and Iso-Seq Analysis on a human dataset. Metrics descriptions are the same as in Table S7.1.

**Table S7.16**: rnaQUAST evaluation results for human dataset SAMN00006579, comparing Scallop-LR, StringTie, and Iso-Seq Analysis

| Metrics | Scallop-LR | StringTie | Iso-Seq |
|---|---|---|---|
| # Transcripts | 20769 | 124624 | 13147 |
| Aligned | 20769 | 124621 | 13147 |
| Uniquely aligned | 20768 | 123959 | 13128 |
| Unaligned | 0 | 3 | 0 |
| Misassemblies | 1 | 25 | 4 |
| 0-50% assembled isoforms | 1214 | 12066 | 1231 |
| 50-75% assembled isoforms | 1516 | 3755 | 1299 |
| 75-95% assembled isoforms | 2494 | 3507 | 2106 |
| 95-100% assembled isoforms | 6387 | 8960 | 3508 |
| Mean isoform assembly | 0.85 | 0.597 | 0.798 |
| 0-50% matched transcripts | 1882 | 20058 | 848 |
| 50-75% matched transcripts | 2384 | 2921 | 545 |
| 75-95% matched transcripts | 4223 | 3167 | 895 |
| 95-100% matched transcripts | 12105 | 9394 | 9655 |
| Unannotated | 174 | 89030 | 1199 |
| Mean fraction of transcript matched | 0.855 | 0.133 | 0.829 |

This table compares the rnaQUAST evaluation results for Scallop-LR, StringTie, and Iso-Seq Analysis on a human dataset. Metrics descriptions are the same as in Table S7.1.

**Table S7.17**: rnaQUAST evaluation results for human dataset SAMN00006580, comparing Scallop-LR, StringTie, and Iso-Seq Analysis

| Metrics | Scallop-LR | StringTie | Iso-Seq |
|---|---|---|---|
| # Transcripts | 18480 | 152741 | 12817 |
| Aligned | 18480 | 152739 | 12817 |
| Uniquely aligned | 18477 | 152257 | 12795 |
| Unaligned | 0 | 2 | 0 |
| Misassemblies | 1 | 17 | 5 |
| 0-50% assembled isoforms | 1246 | 14212 | 1290 |
| 50-75% assembled isoforms | 1497 | 4284 | 1192 |
| 75-95% assembled isoforms | 2438 | 3834 | 2001 |
| 95-100% assembled isoforms | 5809 | 9259 | 3276 |
| Mean isoform assembly | 0.842 | 0.578 | 0.788 |
| 0-50% matched transcripts | 1688 | 24592 | 1023 |
| 50-75% matched transcripts | 1898 | 3141 | 427 |
| 75-95% matched transcripts | 3507 | 2930 | 739 |
| 95-100% matched transcripts | 11154 | 8948 | 8562 |
| Unannotated | 230 | 113096 | 2059 |
| Mean fraction of transcript matched | 0.856 | 0.108 | 0.75 |

This table compares the rnaQUAST evaluation results for Scallop-LR, StringTie, and Iso-Seq Analysis on a human dataset. Metrics descriptions are the same as in Table S7.1.

**Table S7.18**: rnaQUAST evaluation results for human dataset SAMN00006581, comparing Scallop-LR, StringTie, and Iso-Seq Analysis

| Metrics | Scallop-LR | StringTie | Iso-Seq |
|---|---|---|---|
| # Transcripts | 20476 | 125563 | 12512 |
| Aligned | 20476 | 125556 | 12512 |
| Uniquely aligned | 20475 | 125049 | 12495 |
| Unaligned | 0 | 7 | 0 |
| Misassemblies | 1 | 17 | 2 |
| 0-50% assembled isoforms | 1279 | 11940 | 1195 |
| 50-75% assembled isoforms | 1582 | 3652 | 1243 |
| 75-95% assembled isoforms | 2452 | 3340 | 2128 |
| 95-100% assembled isoforms | 6273 | 8602 | 3409 |
| Mean isoform assembly | 0.844 | 0.592 | 0.799 |
| 0-50% matched transcripts | 2036 | 19464 | 878 |
| 50-75% matched transcripts | 2213 | 2901 | 417 |
| 75-95% matched transcripts | 4104 | 2967 | 812 |
| 95-100% matched transcripts | 11946 | 9341 | 9311 |
| Unannotated | 176 | 90846 | 1091 |
| Mean fraction of transcript matched | 0.852 | 0.13 | 0.832 |

This table compares the rnaQUAST evaluation results for Scallop-LR, StringTie, and Iso-Seq Analysis on a human dataset. Metrics descriptions are the same as in Table S7.1.

**Table S8.1**: rnaQUAST evaluation results for mouse dataset SAMEA3374575, comparing Scallop-LR, StringTie, and Iso-Seq Analysis

| Metrics | Scallop-LR | StringTie | Iso-Seq |
|---|---|---|---|
| # Transcripts | 6890 | 12610 | 4017 |
| Aligned | 6890 | 12610 | 4017 |
| Uniquely aligned | 6886 | 12528 | 4013 |
| Unaligned | 0 | 0 | 0 |
| Misassemblies | 4 | 1 | 0 |
| 0-50% assembled isoforms | 703 | 2605 | 606 |
| 50-75% assembled isoforms | 761 | 1258 | 520 |
| 75-95% assembled isoforms | 1194 | 1597 | 1003 |
| 95-100% assembled isoforms | 2451 | 4101 | 1248 |
| Mean isoform assembly | 0.818 | 0.729 | 0.779 |
| 0-50% matched transcripts | 243 | 765 | 41 |
| 50-75% matched transcripts | 369 | 899 | 155 |
| 75-95% matched transcripts | 920 | 1738 | 294 |
| 95-100% matched transcripts | 5323 | 8108 | 3510 |
| Unannotated | 31 | 1098 | 16 |
| Mean fraction of transcript matched | 0.931 | 0.819 | 0.96 |

This table compares the rnaQUAST evaluation results for Scallop-LR, StringTie, and Iso-Seq Analysis on a mouse dataset. Metrics descriptions are the same as in Table S7.1.

**Table S8.2**: rnaQUAST evaluation results for mouse dataset SAMEA3374576, comparing Scallop-LR, StringTie, and Iso-Seq Analysis

| Metrics | Scallop-LR | StringTie | Iso-Seq |
|---|---|---|---|
| # Transcripts | 6720 | 12207 | 4164 |
| Aligned | 6720 | 12206 | 4164 |
| Uniquely aligned | 6715 | 12104 | 4163 |
| Unaligned | 0 | 1 | 0 |
| Misassemblies | 5 | 0 | 1 |
| 0-50% assembled isoforms | 699 | 2406 | 616 |
| 50-75% assembled isoforms | 706 | 1245 | 585 |
| 75-95% assembled isoforms | 1155 | 1632 | 975 |
| 95-100% assembled isoforms | 2563 | 4194 | 1385 |
| Mean isoform assembly | 0.825 | 0.743 | 0.783 |
| 0-50% matched transcripts | 176 | 700 | 48 |
| 50-75% matched transcripts | 275 | 815 | 71 |
| 75-95% matched transcripts | 899 | 1657 | 279 |
| 95-100% matched transcripts | 5334 | 7967 | 3749 |
| Unannotated | 31 | 1067 | 16 |
| Mean fraction of transcript matched | 0.94 | 0.817 | 0.969 |

This table compares the rnaQUAST evaluation results for Scallop-LR, StringTie, and Iso-Seq Analysis on a mouse dataset. Metrics descriptions are the same as in Table S7.1.

**Table S8.3**: rnaQUAST evaluation results for mouse dataset SAMEA3374577, comparing Scallop-LR, StringTie, and Iso-Seq Analysis

| Metrics | Scallop-LR | StringTie | Iso-Seq |
|---|---|---|---|
| # Transcripts | 5972 | 11247 | 3699 |
| Aligned | 5972 | 11247 | 3699 |
| Uniquely aligned | 5967 | 11193 | 3695 |
| Unaligned | 0 | 0 | 0 |
| Misassemblies | 4 | 1 | 1 |
| 0-50% assembled isoforms | 653 | 2250 | 554 |
| 50-75% assembled isoforms | 662 | 1169 | 505 |
| 75-95% assembled isoforms | 1078 | 1531 | 908 |
| 95-100% assembled isoforms | 2343 | 3931 | 1222 |
| Mean isoform assembly | 0.823 | 0.744 | 0.784 |
| 0-50% matched transcripts | 180 | 667 | 60 |
| 50-75% matched transcripts | 236 | 765 | 83 |
| 75-95% matched transcripts | 732 | 1454 | 230 |
| 95-100% matched transcripts | 4796 | 7442 | 3305 |
| Unannotated | 23 | 918 | 20 |
| Mean fraction of transcript matched | 0.942 | 0.826 | 0.963 |

This table compares the rnaQUAST evaluation results for Scallop-LR, StringTie, and Iso-Seq Analysis on a mouse dataset. Metrics descriptions are the same as in Table S7.1.

**Table S8.4**: rnaQUAST evaluation results for mouse dataset SAMEA3374578, comparing Scallop-LR, StringTie, and Iso-Seq Analysis

| Metrics | Scallop-LR | StringTie | Iso-Seq |
|---|---|---|---|
| # Transcripts | 5783 | 10979 | 3492 |
| Aligned | 5783 | 10979 | 3492 |
| Uniquely aligned | 5778 | 10925 | 3491 |
| Unaligned | 0 | 0 | 0 |
| Misassemblies | 5 | 0 | 0 |
| 0-50% assembled isoforms | 625 | 2378 | 543 |
| 50-75% assembled isoforms | 604 | 1166 | 466 |
| 75-95% assembled isoforms | 989 | 1458 | 860 |
| 95-100% assembled isoforms | 2169 | 3674 | 1154 |
| Mean isoform assembly | 0.82 | 0.726 | 0.782 |
| 0-50% matched transcripts | 160 | 696 | 23 |
| 50-75% matched transcripts | 239 | 727 | 57 |
| 75-95% matched transcripts | 842 | 1550 | 210 |
| 95-100% matched transcripts | 4503 | 7184 | 3184 |
| Unannotated | 34 | 822 | 18 |
| Mean fraction of transcript matched | 0.936 | 0.83 | 0.973 |

This table compares the rnaQUAST evaluation results for Scallop-LR, StringTie, and Iso-Seq Analysis on a mouse dataset. Metrics descriptions are the same as in Table S7.1.

**Table S8.5**: rnaQUAST evaluation results for mouse dataset SAMEA3374579, comparing Scallop-LR, StringTie, and Iso-Seq Analysis

| Metrics | Scallop-LR | StringTie | Iso-Seq |
|---|---|---|---|
| # Transcripts | 8052 | 16520 | 4609 |
| Aligned | 8052 | 16520 | 4609 |
| Uniquely aligned | 8052 | 16414 | 4600 |
| Unaligned | 0 | 0 | 0 |
| Misassemblies | 0 | 0 | 0 |
| 0-50% assembled isoforms | 949 | 3648 | 835 |
| 50-75% assembled isoforms | 1006 | 1687 | 650 |
| 75-95% assembled isoforms | 1433 | 1994 | 1111 |
| 95-100% assembled isoforms | 3104 | 5197 | 1502 |
| Mean isoform assembly | 0.812 | 0.714 | 0.763 |
| 0-50% matched transcripts | 155 | 1046 | 26 |
| 50-75% matched transcripts | 288 | 1119 | 82 |
| 75-95% matched transcripts | 989 | 2223 | 268 |
| 95-100% matched transcripts | 6598 | 10605 | 4213 |
| Unannotated | 22 | 1526 | 19 |
| Mean fraction of transcript matched | 0.952 | 0.813 | 0.972 |

This table compares the rnaQUAST evaluation results for Scallop-LR, StringTie, and Iso-Seq Analysis on a mouse dataset. Metrics descriptions are the same as in Table S7.1.

**Table S8.6**: rnaQUAST evaluation results for mouse dataset SAMEA3374580, comparing Scallop-LR, StringTie, and Iso-Seq Analysis

| Metrics | Scallop-LR | StringTie | Iso-Seq |
|---|---|---|---|
| # Transcripts | 8206 | 18482 | 4679 |
| Aligned | 8206 | 18480 | 4679 |
| Uniquely aligned | 8206 | 18301 | 4670 |
| Unaligned | 0 | 2 | 0 |
| Misassemblies | 0 | 0 | 1 |
| 0-50% assembled isoforms | 1057 | 4217 | 1028 |
| 50-75% assembled isoforms | 1016 | 1811 | 627 |
| 75-95% assembled isoforms | 1491 | 1961 | 1047 |
| 95-100% assembled isoforms | 3058 | 5173 | 1408 |
| Mean isoform assembly | 0.802 | 0.692 | 0.729 |
| 0-50% matched transcripts | 149 | 1199 | 42 |
| 50-75% matched transcripts | 281 | 1196 | 57 |
| 75-95% matched transcripts | 960 | 2229 | 257 |
| 95-100% matched transcripts | 6794 | 11398 | 4297 |
| Unannotated | 22 | 2457 | 22 |
| Mean fraction of transcript matched | 0.955 | 0.771 | 0.972 |

This table compares the rnaQUAST evaluation results for Scallop-LR, StringTie, and Iso-Seq Analysis on a mouse dataset. Metrics descriptions are the same as in Table S7.1.

**Table S8.7**: rnaQUAST evaluation results for mouse dataset SAMEA3374581, comparing Scallop-LR, StringTie, and Iso-Seq Analysis

| Metrics | Scallop-LR | StringTie | Iso-Seq |
|---|---|---|---|
| # Transcripts | 9851 | 20349 | 5765 |
| Aligned | 9851 | 20348 | 5765 |
| Uniquely aligned | 9848 | 20157 | 5759 |
| Unaligned | 0 | 1 | 0 |
| Misassemblies | 2 | 2 | 0 |
| 0-50% assembled isoforms | 1199 | 4489 | 1252 |
| 50-75% assembled isoforms | 1137 | 1932 | 743 |
| 75-95% assembled isoforms | 1579 | 2038 | 1204 |
| 95-100% assembled isoforms | 3680 | 5704 | 1695 |
| Mean isoform assembly | 0.808 | 0.696 | 0.728 |
| 0-50% matched transcripts | 221 | 1312 | 57 |
| 50-75% matched transcripts | 418 | 1347 | 108 |
| 75-95% matched transcripts | 1373 | 2690 | 384 |
| 95-100% matched transcripts | 7792 | 12367 | 5166 |
| Unannotated | 44 | 2629 | 50 |
| Mean fraction of transcript matched | 0.944 | 0.772 | 0.964 |

This table compares the rnaQUAST evaluation results for Scallop-LR, StringTie, and Iso-Seq Analysis on a mouse dataset. Metrics descriptions are the same as in Table S7.1.

**Table S8.8**: rnaQUAST evaluation results for mouse dataset SAMEA3374582, comparing Scallop-LR, StringTie, and Iso-Seq Analysis

| Metrics | Scallop-LR | StringTie | Iso-Seq |
|---|---|---|---|
| # Transcripts | 8071 | 18712 | 4748 |
| Aligned | 8071 | 18712 | 4748 |
| Uniquely aligned | 8069 | 18543 | 4740 |
| Unaligned | 0 | 0 | 0 |
| Misassemblies | 1 | 1 | 0 |
| 0-50% assembled isoforms | 1115 | 4837 | 1344 |
| 50-75% assembled isoforms | 1081 | 1980 | 619 |
| 75-95% assembled isoforms | 1335 | 1892 | 919 |
| 95-100% assembled isoforms | 2718 | 4581 | 1187 |
| Mean isoform assembly | 0.784 | 0.657 | 0.674 |
| 0-50% matched transcripts | 98 | 1091 | 36 |
| 50-75% matched transcripts | 243 | 1166 | 43 |
| 75-95% matched transcripts | 905 | 2275 | 170 |
| 95-100% matched transcripts | 6806 | 11737 | 4444 |
| Unannotated | 17 | 2439 | 55 |
| Mean fraction of transcript matched | 0.962 | 0.779 | 0.97 |

This table compares the rnaQUAST evaluation results for Scallop-LR, StringTie, and Iso-Seq Analysis on a mouse dataset. Metrics descriptions are the same as in Table S7.1.

**Table S9**: SRA Information for the 26 Datasets Used in this Study

| Dataset | BioSample | SRA Study | Organism | Year | Instrument |
|---|---|---|---|---|---|
| 1 | SAMN00001694 | ERP015321 | *Homo sapiens* | 2016 | RS |
| 2 | SAMN00001695 | ERP015321 | *Homo sapiens* | 2016 | RS |
| 3 | SAMN00001696 | ERP015321 | *Homo sapiens* | 2016 | RS |
| 4 | SAMN00006465 | ERP015321 | *Homo sapiens* | 2016 | RS |
| 5 | SAMN00006466 | ERP015321 | *Homo sapiens* | 2016 | RS |
| 6 | SAMN00006467 | ERP015321 | *Homo sapiens* | 2016 | RS |
| 7 | SAMN00006579 | ERP015321 | *Homo sapiens* | 2016 | RS |
| 8 | SAMN00006580 | ERP015321 | *Homo sapiens* | 2016 | RS |
| 9 | SAMN00006581 | ERP015321 | *Homo sapiens* | 2016 | RS |
| 10 | SAMN08182059 | SRP126849 | *Homo sapiens* | 2017 | RS II |
| 11 | SAMN08182060 | SRP126849 | *Homo sapiens* | 2017 | RS II |
| 12 | SAMN04563763 | SRP071928 | *Homo sapiens* | 2016 | RS II |
| 13 | SAMN07611993 | SRP098984 | *Homo sapiens* | 2018 | RS II |
| 14 | SAMN04169050 | SRP068953 | *Homo sapiens* | 2016 | RS II |
| 15 | SAMN04251426.1 | SRP065930 | *Homo sapiens* | 2016 | RS II |
| 16 | SAMN04251426.2 | SRP065930 | *Homo sapiens* | 2016 | RS II |
| 17 | SAMN04251426.3 | SRP065930 | *Homo sapiens* | 2016 | RS II |
| 18 | SAMN04251426.4 | SRP065930 | *Homo sapiens* | 2016 | RS II |
| 19 | SAMEA3374575 | ERP010189 | *Mus musculus* | 2015 | RS |
| 20 | SAMEA3374576 | ERP010189 | *Mus musculus* | 2015 | RS |
| 21 | SAMEA3374577 | ERP010189 | *Mus musculus* | 2015 | RS |
| 22 | SAMEA3374578 | ERP010189 | *Mus musculus* | 2015 | RS |
| 23 | SAMEA3374579 | ERP010189 | *Mus musculus* | 2015 | RS |
| 24 | SAMEA3374580 | ERP010189 | *Mus musculus* | 2015 | RS |
| 25 | SAMEA3374581 | ERP010189 | *Mus musculus* | 2015 | RS |
| 26 | SAMEA3374582 | ERP010189 | *Mus musculus* | 2015 | RS |

This table summarizes the 26 datasets used in this paper. 18 datasets are human and eight datasets are mouse. The data were downloaded from the corresponding SRA Study. The multiple SRA Runs for each BioSample under the corresponding SRA Study were extracted, processed, and then merged into a large dataset.
